# Acoustic Force Spectroscopy Reveals Subtle Differences in Cellulose Unbinding Behavior of Carbohydrate-Binding Modules

**DOI:** 10.1101/2021.09.20.461102

**Authors:** Markus Hackl, Edward V. Contrada, Jonathan E. Ash, Atharv Kulkarni, Jinho Yoon, Hyeon-Yeol Cho, Ki-Bum Lee, John M. Yarbrough, Shishir P. S. Chundawat

**Affiliations:** Department of Chemical and Biochemical Engineering, Rutgers, The State University of New Jersey, Piscataway New Jersey, USA; Department of Chemistry and Chemical Biology, Rutgers, The State University of New Jersey, Piscataway New Jersey, USA; Biosciences Center, National Renewable Energy Laboratory, Golden, Colorado, USA

**Author notes:** Corresponding author: Shishir P. S. Chundawat **Email:**. **Author Contributions:** M.H. and S.P.S.C. designed research; M.H., E.C., J.E.A, A.K., J.Y., H.C., K.L. and J.M.Y. performed experiments; M.H., E.C. and J.E.A. analyzed the data; M.H. and S.P.S.C wrote the manuscript with inputs from all authors.

**Keywords:** Carbohydrate-Binding Module, Nanocellulose, Biofuels, Single-Molecule Force Spectroscopy, Acoustic Force Spectroscopy

## Abstract

To rationally engineer more efficient cellulolytic enzymes for cellulosic biomass deconstruction into sugars for biofuels production, it is necessary to better understand the complex enzyme-substrate interfacial interactions. Carbohydrate binding modules (CBM) are often associated with microbial surface-tethered cellulosomal or freely secreted cellulase enzymes to increase substrate accessibility. However, it is not well known how CBM recognize, bind, and dissociate from polysaccharide surfaces to facilitate efficient cellulolytic activity due to the lack of mechanistic understanding of CBM-substrate interactions. Our work outlines a general approach to methodically study the unbinding behavior of CBMs from model polysaccharide surfaces using single-molecule force spectroscopy. Here, we apply acoustic force spectroscopy (AFS) to probe a *Clostridium thermocellum* cellulosomal scaffoldin protein (CBM3a) and measure its dissociation from nanocellulose surfaces at physiologically relevant, low force loading rates. An automated microfluidic setup and methodology for uniform deposition of insoluble polysaccharides on the AFS chip surfaces is demonstrated. The rupture forces of wild-type CBM3a, and its Y67A mutant, unbinding from nanocellulose surface suggests distinct CBM binding conformations that can also explain the improved cellulolytic activity of cellulase tethered to CBM. Applying established dynamic force spectroscopy theory, the single-molecule unbinding rate at zero force is extrapolated and found to agree well with bulk equilibrium unbinding rates estimated independently using quartz crystal microbalance with dissipation monitoring. However, our results highlight the limitations of applying classical theory to explain the highly multivalent CBM-cellulose interactions seen at higher cellulose-CBM bond rupture forces (>15pN).

**Significance Statement:** Cellulases are multi-modular enzymes produced by numerous microbes that catalyze cellulose hydrolysis into glucose. These enzymes play an important role in global carbon cycling as well as cellulosic biofuels production. CBMs are essential components of cellulolytic enzymes involved in facilitating hydrolysis of polysaccharides by tethered catalytic domains (CD). The subtle interplay between CBM binding and CD activity is poorly understood particularly for heterogeneous reactions at solid-liquid interfaces. Here, we report a highly multiplexed single-molecule force spectroscopy method to study CBM dissociation from cellulose to infer the molecular mechanism governing substrate recognition and dissociation. This approach can be broadly applied to study multivalent protein-polysaccharide binding interactions relevant to other carbohydrates such as starch, chitin, or hyaluronan to engineer efficient biocatalysts.

## Introduction

Carbohydrate-based biopolymers are abundant throughout all forms of life and play a major part in biomolecular recognition processes that has fundamental scientific and applied technological relevance. For example, the adsorption of enzymes secreted by cellulolytic microbes to carbohydrate polymers, like cellulose, is important in deconstructing lignocellulosic biomass to fermentable sugars for biofuel production (1, 2). Carbohydrate-Active enZymes (CAZymes) such as processive cellulases often consist of two or more domains called carbohydrate-binding modules (CBM) and catalytic domains (CD), which are responsible for the recognition/binding and breakdown of the substrate, respectively (3). On the other hand, cellulosomes are larger multidomain enzymes where CDs are assembled on a scaffolding domain decorated with CBMs and specific linker domains as shown in **Figure 1-A** (4). Cellulosomes adapt to the substrate topology and display a “sit-and-dig” mechanism where the cellulosome degrades individual cellulose crystals without dissociating from the substrate (5–7). This mode of action contrasts processive cellulases such as *Trichoderma reesei* Cel7A, which displays a “slide-and-peel” mechanism and frequently dissociate from the substrate (3, 8–10).

**Figure 1.**
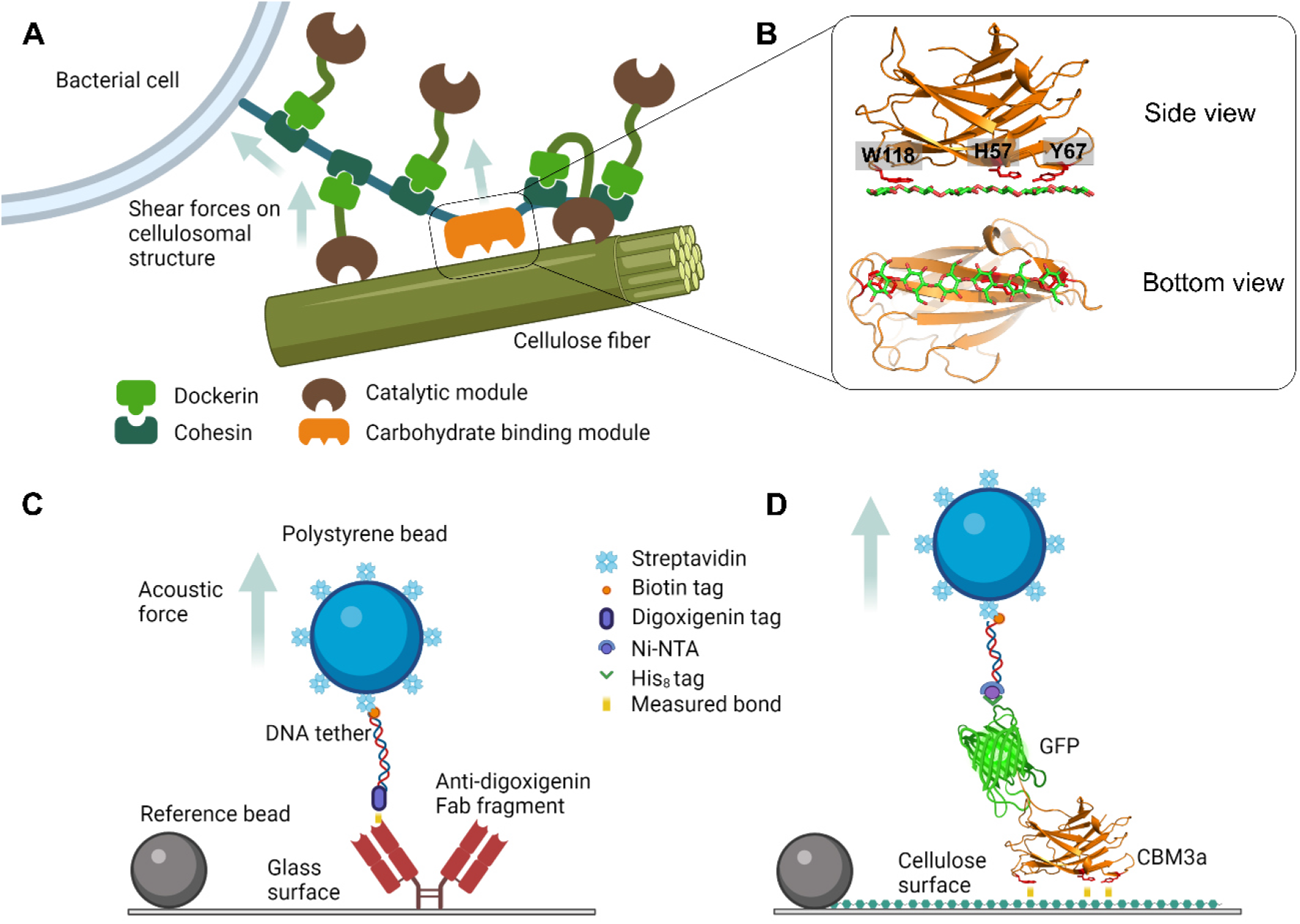
Schematic of a generic cellulosome and acoustic force spectroscopy experimental setups to characterize single-molecule model protein-ligand and CBM3a-polysaccharide unbinding forces (not drawn to scale). A) Generic bacterial cell surface anchored cellulosome is shown adhering to a single cellulose fiber. The carbohydrate-binding module (CBM) binds to cellulose and directs the catalytic domains to the cellulose surface. Shear forces due to the substrate or cell movement are exerted on the cellulosome scaffold. B) Side and bottom view of CBM3a structure with key aromatic residues involved in binding to cellulose (PDB: 1nbc). The aromatic residues (W118, H57, and Y67) form a flat binding surface complementary to the cellulose surface. C) Schematic outlining the measurement of the unbinding force of model digoxigenin (DIG) ligand from surface-bound anti-DIG antibody to validate the bead preparation method as well as analysis procedure of AFS traces. D) Schematic outlining the measurement of the unbinding force of His-GFP tagged CBM3a from a nanocrystalline cellulose surface using the AFS assay.

CBMs can be grouped into type-A, B, or C categories based on relevant structural-functional relationships. Both *Tr*Cel7A and the cellulosome from *Clostridium thermocellum* possess a type-A CBM with a similar architecture of the cellulose binding site. Type-A CBMs preferably bind to insoluble and highly crystalline cellulose, forming a flat, platform-like binding surface mostly lined with aromatic residues, complementary to the flat planar structure of the crystalline substrate (11). As such, CBM1 from *Tr*Cel7A exhibits 3 tyrosine residues at the 5, 31, and 32 positions (12), whereas the 3 aromatic residues on the binding surface of *Ct*CBM3a are H57, Y67, and W118 respectively (13) as shown in **Figure 1-B**. Although mutations of the aromatic residues of CBM3a to alanine can reduce the apparent bulk-ensemble binding affinity to native crystalline cellulose, the enzymatic activity of endocellulases fused to those mutants increased by 20-70% compared to the wild-type (14). Altering enzyme binding affinity to cellulosic substrates is being explored as a strategy to engineer more efficient cellulases (15, 16). However, engineering highly active cellulases, cellulosomes, and associated cellulolytic microbes still present challenges due to the inadequate understanding of the complex interplay between CD and CBM as well as the multivalent nature of the CBM-carbohydrate binding interactions.

Traditionally, CBM and cellulase adsorption is characterized by bulk ensemble-based methods such as solid-phase depletion (17, 18), quartz crystal microbalance with dissipation (QCM-D) (19), and isothermal titration calorimetry (ITC) (20, 21). But these methods rely on simplified models to illustrate binding interactions that do not reflect the underlying molecular mechanism of protein binding to highly multivalent carbohydrate ligands such as cellulose. Techniques like singlemolecule fluorescence (10, 22) and force spectroscopy (23) have greatly contributed to our understanding of molecular processes relevant to cellulose degradation. In particular, atomic force microscopy (AFM) has been used previously to characterize CBM desorption from cellulose on the single-molecule level (24, 25). Examples include the identification of binding sites (26), distinguishing specific from non-specific binding (27), and determining the zero-force unbinding rate using Bell’s model (28). Although AFM measures force and distance with pN and nm resolution, the determination of unbinding forces occurs far from equilibrium due to the relatively high loading rates inherent to the AFM instrument, potentially obscuring multimodal unbinding behavior seen at physiologically relevant conditions. Alternatively, optical tweezers (OT) have been used to study CBM unbinding at lower loading rates and force clamp mode (29), and the results suggest a more complex unbinding behavior where the bond-lifetime data does not follow a single exponential decay function as suggested by AFM studies (28).

In contrast to AFM, acoustic force spectroscopy (AFS) is a new technique that enables the application of low loading rates comparable to optical tweezers while maintaining a higher throughput (30, 31). Similar to OT, the molecule of interest is attached to a micrometer-sized bead *via* a double-stranded DNA tether. However, forces on the bead are exerted by acoustic standing waves. Most commonly, streptavidin-coated beads are used to connect biotinylated DNA tethers (32) due to high specificity and binding strength (33, 34). On the other end of the tether, the protein of interest is either covalently linked through a thiol-maleimide crosslink (23) or tethered non-covalently *via* the histidine tag to Anti-his antibodies, which in turn are covalently linked to aminated DNA tethers (29). The histidine tag of proteins was used previously to non-covalently link to DNA (35–38) and directly attaching proteins to NTA-modified AFM tips (39). It was demonstrated that the His-Ni-NTA bond is stable enough (~120pN at 400pN/s) to facilitate single-molecule force spectroscopy (SMFS) experiments of the tethered protein (40–43) thus allowing AFM-based studies measuring unbinding forces of CBM3a using Ni-NTA (25),(27).

Here, we combine the tethering methods by directly synthesizing a linear dsDNA tether with biotin on one end to attach it to a micron-sized bead, and Ni-NTA on the other end to tether any His-tagged protein. This setup allows SMFS of the His-tagged protein with its polysaccharide ligand deposited onto the AFS microfluidic channel surface to enable high-throughput assays. Such tethers can be used in other tethered bead setups such as optical or magnetic tweezers, highlighting the modularity and versatility of our approach. Furthermore, digoxigenin (DIG) tethers instead of NTA were generated with the same procedure to validate the bead preparation method as well as analysis of the recorded position and force-distance traces using AFS. A schematic of the protein-ligand systems studied to measure bond rupture forces is shown in **Figure 1-C** and D. Furthermore, an automated method for the deposition of nanocrystalline cellulose (NCC) inside the AFS chip is developed. The unbinding forces of *C. thermocellum* CBM3a-wt (WT) and its Y67A mutant are measured at fixed, low loading rates. The unbinding forces of the wild-type have been previously characterized by AFM (25–28). It was shown previously that the Y67A mutation reduces CBM binding affinity by several orders of magnitude while improving tethered CD activity for reasons not clear (14). Furthermore, the unbinding behavior of the wild-type and mutant CBM3a measured using SMFS at physiologically relevant conditions has not been reported. We identified a clear difference in the rupture force distribution observed between WT and Y67A mutant at low loading rates. Lastly, while the extracted unbinding rate (*k_off_*) from our AFS results agrees with bulk ensemble QCM-D results, the classical SMFS model is unable to accurately capture the multivalent protein-polysaccharide binding interactions particularly at higher rupture forces.

## Results

### Deposition and characterization of nanocellulose inside the AFS chip

Sulfuric acid-derived nanocrystalline cellulose (NCC) was used to generate the cellulose model film in this study. The formation of an NCC film inside the AFS chip was accomplished by a multilayer deposition process (44) where poly-L-Lysine (PLL) and NCC were alternatingly deposited using an automated microfluidic control system. **Figure 2-A** and B show the flowchart and process flow diagram of the process, and a detailed description can be found in the methods section. Green fluorescent protein (GFP) tagged CBM3a was expressed as described previously (14) and used to characterize the cellulose film deposited on the AFS chip. **Figure 2-C** shows a representative fluorescence image of the NCC-modified AFS chip labeled with GFP-CBM3a. The arrow indicates an area where a bubble was stuck during the NCC deposition process. Slightly lower amounts of NCC were deposited in that area, resulting in lower fluorescence. The rest of the flow channel displays a uniform fluorescence, indicating that NCC is deposited evenly across the channel. The average fluorescence intensity of the bare glass and PLL treated chips surfaces is 14 ± 1 a.u. (mean ± SEM) and 12 ± 3 a.u., respectively, whereas the NCC treated chips show a fluorescence intensity of 136 ± 35 a.u. The deposition of a single layer of NCC onto a PLL treated surface resulted in a fluorescence intensity of 42 ± 24 a.u. Despite significantly higher fluorescence compared to controls, such prepared AFS chips failed to reproducibly provide a consistent response at the single-molecule level even though AFM imaging confirmed the deposition of a uniform layer of NCC (**SI Appendix Fig. S1**). A relationship between the success of a single-molecule experiment and the measured fluorescence intensity was observed, where the likelihood of a successful singlemolecule experiments positively correlated with the measured fluorescence. Hence, a multilayer NCC deposition method was applied to ensure a consistently high fluorescence signal, which in turn resulted in a reliable rupture force measurement of CBMs. Multilayer NCC-functionalized AFS chips, which were subsequently cleaned and imaged as outlined in the methods section, showed a fluorescence intensity of 13±0.5, indicating the removal of nanocrystalline cellulose for reuse of the AFS chips for multiple rounds of experimentation. **Figure 2-D** shows an example surface imaged by AFM, additional AFM images of bare and PLL treated surfaces are found in **SI Appendix Fig. S1**. Similar to spin-coated samples (44, 45), the surface was uniformly covered with nanocrystalline cellulose. AFM image analysis revealed the formation of NCC crystal aggregates at multilayered films. This is reflected by a surface roughness factor (Ra) that is marginally greater than 3 nm compared to less than 2 nm estimated for a single NCC layer.

**Figure 2.**
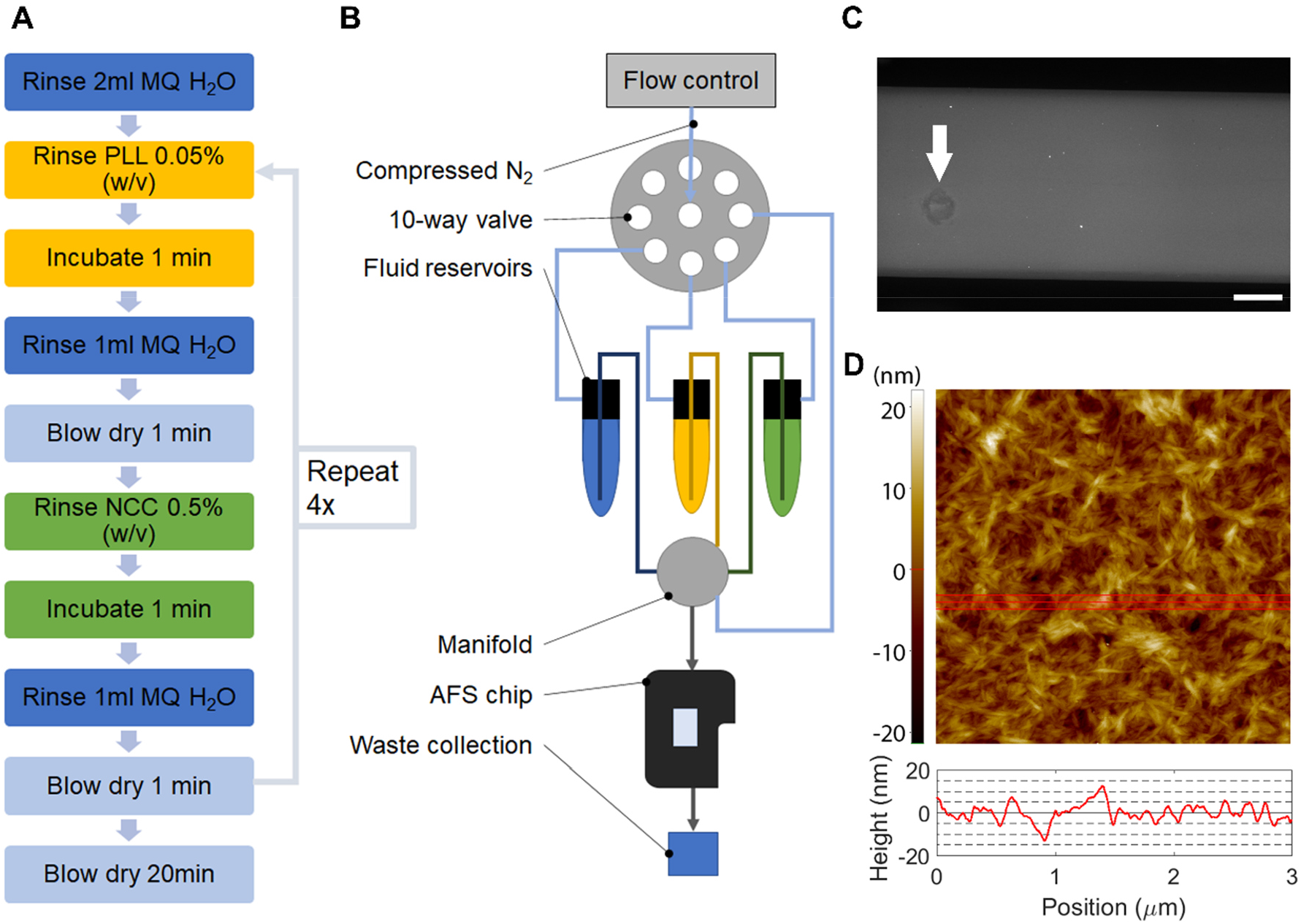
Multilayer deposition of nanocrystalline cellulose (NCC) within the AFS chip enables the characterization of a uniform and reproducible surface. A) Flowchart and B) Process flow diagram of the NCC deposition method, C) Fluorescence image of the NCC modified AFS chip. GFP-CBM3a WT was used to bind to and visualize the deposited NCC film. The arrow indicates a representative area where a bubble was stuck at some point during the NCC deposition, hence a lesser amount of GFP-CBM3a is bound to that area. The scale bar is 500 μm. D) AFM image (3×3 μm) of the NCC film deposited on a glass slide showing a densely covered surface. The red line represents the area used to obtain the average height profile trace shown below. Despite minor aggregation of NCC crystals during layer-by-layer deposition, height differences are less than 20 nm.

### Observation of shortened DNA tethers on NCC surfaces

The tether preparation method and analysis of traces as outlined in the methods section were validated by tethering DNA anchored to the AFS chip surface by anti-digoxigenin antibodies (aDIG). The dimensionless contour length (*l_fc_*) of DNA tethers bound to aDIG during force calibration was 1.1 ± 0.12 (mean ± SD, N=156). This is in the expected range given the particle size distribution of the beads. The average rupture force of the DIG-aDIG complex was determined to be 18.8 ± 7.0 pN at a loading rate of 0.14 ± 0.05 pN/s (**SI Appendix Fig. S2**) and is close to the reported value of 16.6 pN at 0.11 pN/s (30). As shown in **Figure 3-A**, overstretching of the DNA tether was observed at ~65pN, thus confirming the formation of single tethers with the bead preparation method outlined in the methods section. In contrast, the observed dimensionless tether length for NCC-CBM tethered DNA was only 0.83±0.23, indicating a shortening of the tethers by ~25%. However, the force-distance (FD) curves obtained during the linear force ramp follow the extensible worm-like chain (46) or WLC model (**Figure 3-B**), indicating that the tethers are only shortened but not otherwise altered. **Figure 3-C** and D show the scatter plots of the rupture force with *l_fc_* for DIG-aDIG and NCC-CBM3a-wt at 1 pN/s respectively. The best linear fit (red line) is added as a guide. Additional scatter plots of root-mean-square fluctuation (*RMS*) and symmetrical motion (*Sym*) as well as the Pearson and Spearman correlation coefficients can be found in the **SI Appendix Fig. S3** and **Table S1** and **Table S2**. Except for the Pearson for *Sym* and rupture force of Y67A at 0.1 pN/s (p=0.043), no significant correlation (p<0.05) was identified between the measured rupture force and observed length as well as *RMS* and *Sym*. We hypothesize that NCC surface-displayed crystals wrap around and/or interact with the DNA during the incubation step when the DNA is near the surface. The attached NCC crystals are subsequently detached from the NCC surface when the bead is being pulled away from the surface during force calibration but stay bound or interact with the DNA. Non-specifically tethered beads were observed in control experiments with blank Ni-NTA and GFP tagged beads. However, the number of tethered beads was higher by at least 4x for CBM tethered beads. The loss of tethered beads during the flushing step before bead tracking was noted in all cases but was significantly larger for non-CBM tethered beads further indicating weaker, nonspecific binding interactions of the DNA to NCC. The rupture force distribution of only tethers close to the expected length and the entire expected single-molecule tethers are identical as it can be seen in **SI Appendix Fig. S4**, implying that a single CBM-NCC rupture event was measured even though a shortened tether was observed. Assuming that a single CBM was tethered when the FD curve follows the WLC model, the force calibration and rupture force determination were not affected by the shortening of DNA, the data were included in all further analyses.

**Figure 3.**
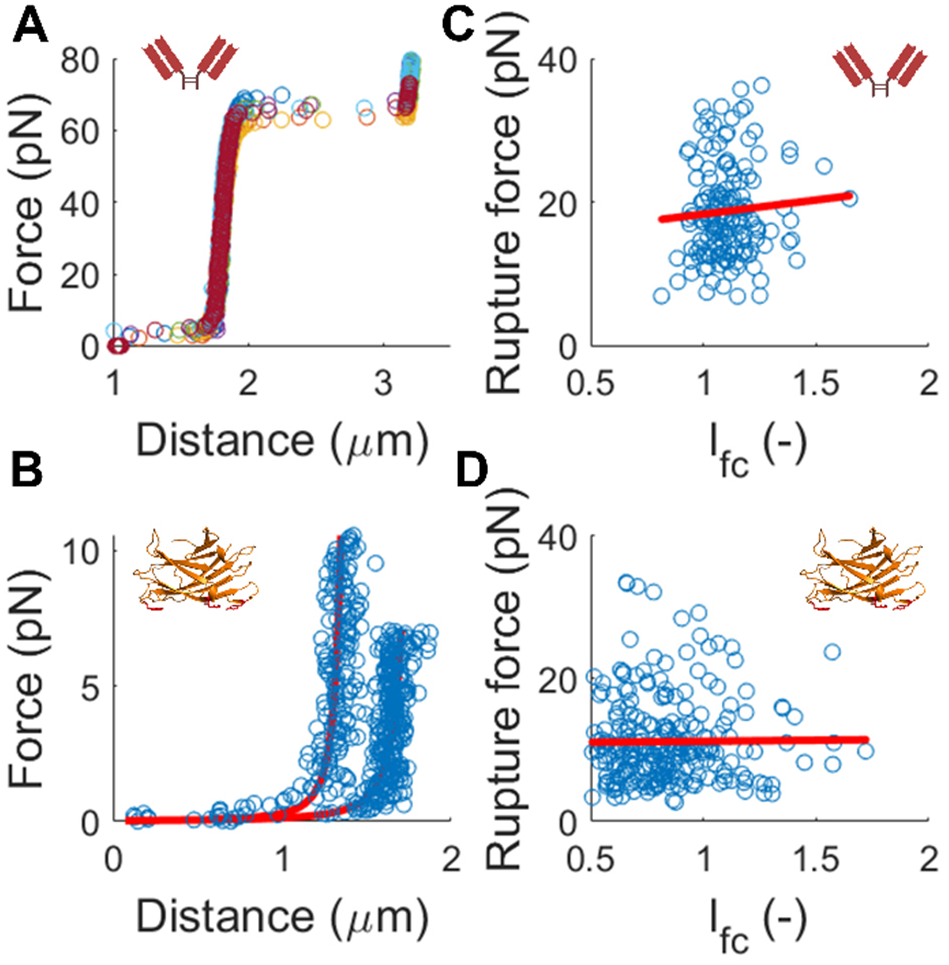
No correlation was observed between tether length and rupture force for DIG-aDIG and CBM3a-cellulose interactions. A) Force-distance (FD) curves of DNA anchored to the chip surface by the DIG-aDIG bond (N=7). The extension at ~65 pN is characteristic for a single DNA tether and indicates overstretching of DNA. B) Example of FD curves for DNA anchored to the chip surface by the NCC-CBM-bond. The red line shows the WLC fit with *l_p_*=42 nm and S=1300 pN. Despite following the WLC model, the tethers show a reduction in length of 25% on average. No overstretching was observed since CBMs detach from the surface well below 65 pN. C) Scatterplot and linear fit (red line) of rupture force and dimensionless length during force calibration (*l_fc_*) for DIG-aDIG (N=156). D) Scatterplot and linear fit (red line) of rupture force and *l_fc_* for CBM3a-wt at 1 pN/s (N=259). No significant correlation is found between the measured rupture force and *l_fc_* (See **SI Appendix Table S1** and **Table S2**).

### Rupture force analysis and application of the Dudko-Hummer-Szabo (DHS) model

The rupture force distributions for CBM3a-WT and Y67A mutant at a loading rate of 1 pN/s and 0.1 pN/s are shown in **Figure 4**. The bin width for the histograms was chosen based on the Freedman-Diaconis rule (47) since the data deviates from a single normal distribution. To capture the apparent multimodal distribution, a double normal distribution was fit to the histogram. The means and standard deviations are summarized in **SI Appendix Table S3**. Although the first mean is similar for wild-type (8.5 pN) and Y67A (7.9 pN) at 1 pN/s, there is a clear single rupture force peak observed for the Y67A mutant, but not for the wild type. This difference is even more pronounced when comparing the rupture force distributions at 0.1 pN/s. Two distinct rupture force peaks were observed for the wild type at 3.5 pN and 7.1 pN, respectively, whereas Y67A showed only one peak at 4.5 pN. All histograms show a “tail” towards larger rupture forces, which is defined by the second normal fit. At 1 pN/s, CBM3a-WT shows a distinct peak at 17.5 pN, followed by a long tail up to 35 pN, whereas no clear second peak was seen but only a tail until 25 pN is observed for Y67A.

**Figure 4.**
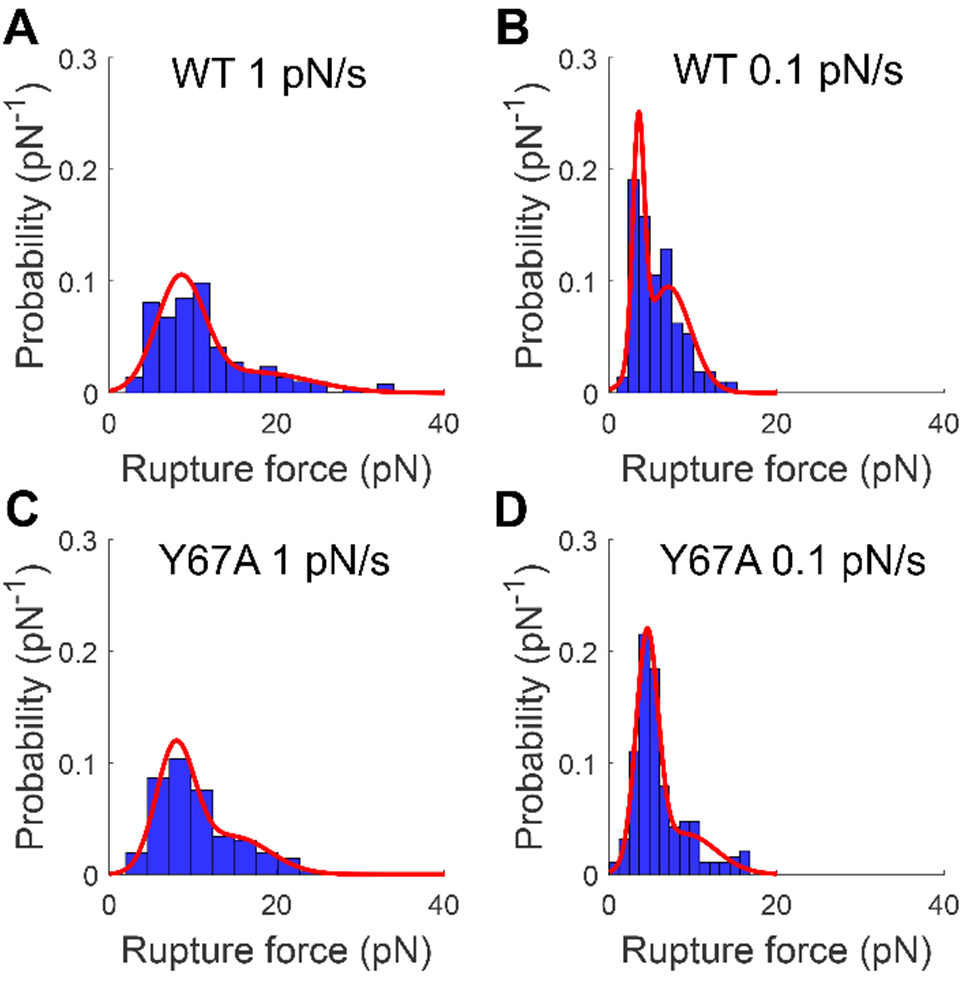
AFS technique highlights distinct multimodal CBM-cellulose rupture forces distribution at lower loading rates. A-B) Obtained rupture force histograms and fit to a double normal distribution for CBM3a WT at a loading rate of 1 pN/s (N=259) and 0.1 pN/s (N=161) respectively. C-D) Rupture force histograms and fit to a double normal distribution for CBM3a Y67A at a loading rate of 1 pN/s (N=138) and 0.1 pN/s (N=159). The fit parameters are summarized in **SI Appendix Table S3**. The tail towards higher rupture forces is observed in all cases, however, CBM3a Y67A displays only a single peak at both loading rates, whereas CBM3a-WT shows no clear single peak, but rather 2 or more rupture force peaks.

**Figure 5-A** and B show the transformation of rupture force histograms to force-dependent bond-lifetime data using Equation 1 (circles) and the fit of Equation 2 (solid lines) of the Dudko-Hummer-Szabo (DHS) model (48) described in the methods section, for WT and Y67A respectively. Data from rupture force histograms obtained at different loading rates should fall on the same master curve for force-dependent bond-lifetimes as predicted by Equation 2 if the unbinding kinetics at constant force follows a single-exponential function (49). Although there is some overlap of bond-lifetimes obtained at 0.1 pN/s and 1 pN/s for both WT and Y67A, the fitted Equation 2 inadequately describes the data for both shape factors *v*. A similar observation of bond-lifetime data not following classical models was made recently for another type-A CBM1 and its Y31A mutant using optical tweezers (29), although the force-dependent bond-lifetime was obtained in force-clamp mode. Surprisingly, no significant difference in the force-dependent bond-lifetime was found between CBM1 and its Y31A mutant for most rupture forces. **Figure 5-C** and D show the rupture force histograms of WT and Y67A and the predicted probability density according to Equation 3. Both shape factors produce a qualitatively similar probability distribution but insufficiently replicate the measured rupture forces. The main reason for discrepancies in bond-lifetime and rupture probability distribution fit is the shape of the underlying rupture force histogram. Both the WT and Y67A rupture force histograms show tailing towards higher rupture forces with no clear peak, which results in almost force-independent bond-lifetimes at higher rupture forces. The multimodal distribution observed for WT at both loading rates results in bond-lifetime data not exactly following a single exponential decay function. As shown in **Table 1**, only *v*=2/3 yields unity for the numerical approximation of ∫ *p*(*f*)*df* over the modelled force range, despite qualitatively similar fits of the bond-lifetime data and probability density for both shape factors. The extrapolated unbinding rates (*k_off_*) at zero force and *v*=2/3 for the WT is 0.0091s^−1^ and approximately twice as high as the *k_off_* for Y67A at 0.0044s^−1^.

**Figure 5.**
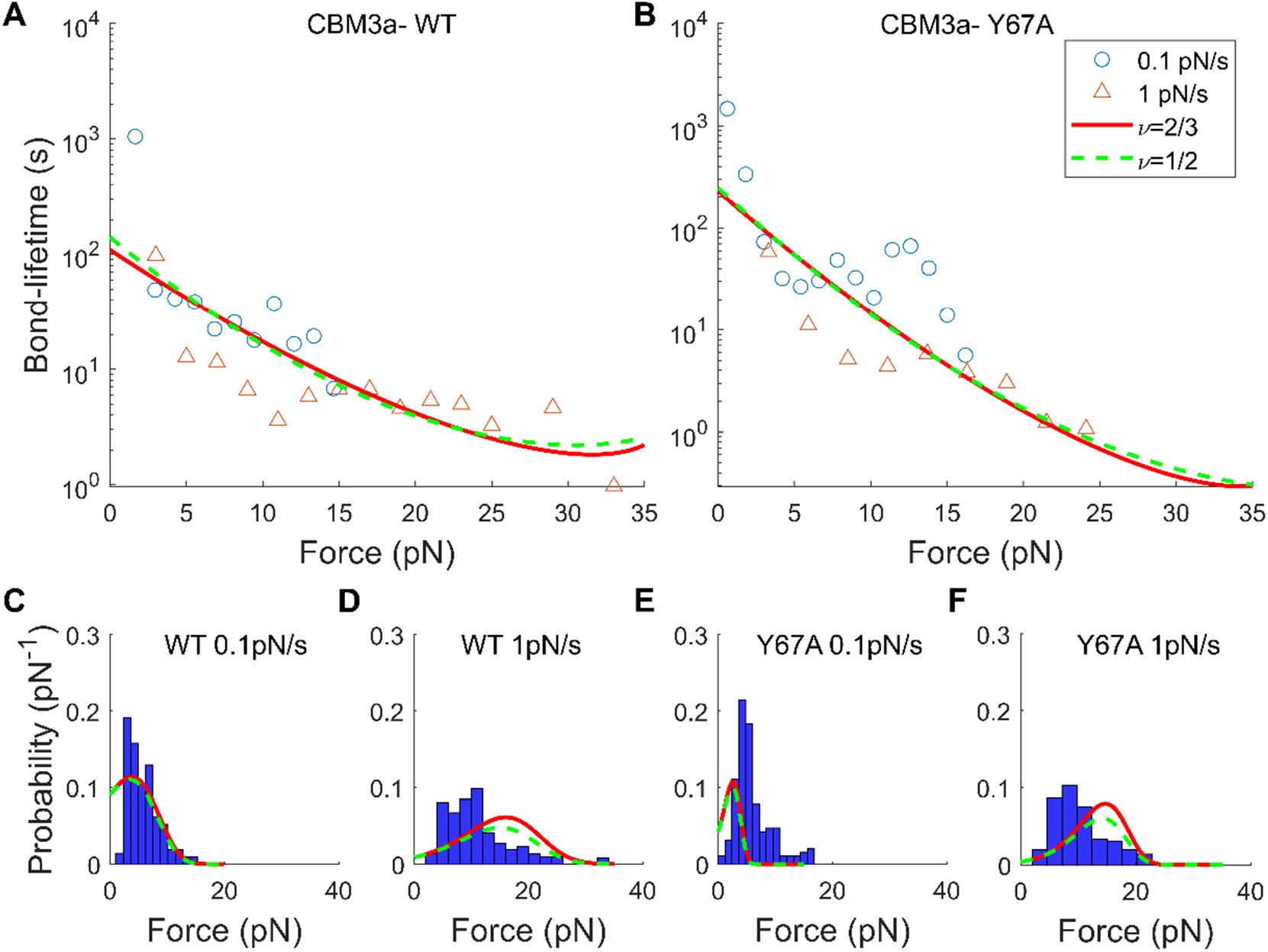
Application of the DHS model to obtained CBM3a-cellulose rupture forces highlights limitations of classical theory to study multivalent protein-polysaccharide unbinding interactions. A, B) Force-dependent bond-lifetime obtained from transforming rupture force distributions at 0.1 pN/s (o) and 1 pN/s (Δ) using Equation 1 for WT and Y67A, respectively. The fit of Equation 2 is shown for *v*=2/3 (red, solid line) and *v*=1/2 (green, dashed line). C-F) Rupture force distributions at 0.1 pN/s and 1 pN/s with the fit of Eq. 3 for WT and Y67A respectively, using the parameters obtained from fitting Equation. 2 to data in A) and B) for *v*=2/3 (red, solid line) and *v*=1/2 (green, dashed line). While both shape factors yield a qualitatively similar fit, only *v*=2/3 results in *∫ p*(*f*)*df* = 1.

**Table 1.** Fit parameters of force-dependent bond lifetimes for CBM3a WT and Y67A mutant, respectively. The integral column refers to the numerical integration of the rupture force probability as described by Equation 3 in the Materials and Methods section for both loading rates.

| | $\nu$<br>(-) | $k_{off}$<br>(s<sup>-1</sup>) | $x^\ddagger$<br>(nm) | $\Delta G^\ddagger$<br>(k<sub>B</sub>T) | $\int p(f)df$ at 0.1 pN/s [1 pN/s]<br>(-) |
| --- | --- | --- | --- | --- | --- |
| **WT** | 2/3 | 0.0091 | 0.88 | 5.4 | 1.0 [1.0] |
|  | 1/2 | 0.0071 | 1.24 | 5.9 | 0.95 [0.80] |
| **Y67A** | 2/3 | 0.0044 | 1.12 | 8.1 | 1.0 [1.0] |
|  | 1/2 | 0.0041 | 1.22 | 8.7 | 0.93 [0.79] |

Table 1 summarizes the fit parameters from Equation 2 as well as the numerical approximation of ∫ *p*(*f*)*df* for both loading rates. The transition state distance, x^‡^, is 0.88 nm and 1.12 nm for CBM3a WT and Y67A, respectively, and in agreement with a transition state distance based on Bell’s model of 0.63 nm for CBM3a WT (28). The apparent free energy of activation, Δ*G*^‡^, is 5.4 *k_B_T* and 8.1 *k_B_T* for CBM3a-wt and Y67A respectively, and contrasts 45.3 k_B_T previously reported (28). Both x^‡^ and Δ*G*^‡^ are similar for the wild-type and mutant, indicating a similar unbinding pathway.

### Bulk ensemble CBM3a-nanocellulose off-rate qualitatively agrees with SMFS result

QCM-D experiments using hydrochloric acid-derived NCC as substrate reported a 1.4-fold increase in the off-rate for the Y67A mutant compared to the WT (14). However, using sulfuric acid-derived NCC, our QCM-D analysis using a classical one-site binding site adsorption model yields a *k_off_* of 26.8 ± 2.4 ×10^−5^ s^−1^ (mean ± SD, n=2) and 19.7 ± 1.2 ×10^−5^ s^−1^ for WT and Y67A respectively. This result supports the findings from AFS experiments that the WT unbinds more frequently, although the absolute values differ between AFS and QCM-D. In contrast, the number of available binding sites determined by QCM-D reduced from 306 ± 41 ×10^12^ molecules to 177 ± 43 ×10^12^ molecules between WT and Y67A, respectively. The unbinding rate for CBM3a WT, from sulfuric acid-derived microfibrils isolated from poplar, extracted from SMFS rupture force data using the Bell’s model was estimated to be 0.0089 s^−1^ (28) and is close to the value obtained in our study. However, both the Bell and DHS models assume a one-dimensional unbinding pathway, which may not represent the underlying molecular interactions based on the multimodal rupture force distributions measured in this study, as well as evidence of different CBM binding orientations to crystalline cellulose that give rise to multiple non-equivalent binding sites (50),(29).

## Discussion

We established a layer-by-layer deposition method for the immobilization of nanocrystalline cellulose onto microfluidic chip surfaces and determined single-molecule CBM-cellulose unbinding forces at varying loading rates using AFS. Any soluble or insoluble polysaccharide substrate that can be spin-coated on glass surfaces and is small enough not to clog the flow channel, can be readily immobilized using our proposed approach. Examples include the immobilization of regenerated cellulose, cellulose microfibrils, or chitin nanocrystals (24, 51, 52). Cellulose nanocrystals offer an especially promising platform for further chemical modifications (53), (54) either pre-or post-immobilization to fine-tune protein adsorption (55), allowing the application of SMFS to a wide range of applications. Furthermore, a robust method for preparing tethered beads utilizing the commonly used biotin-streptavidin and His-Ni-NTA interactions is presented. Histidine tags are widely used to purify heterologously expressed proteins, thus the proposed one-step tether synthesis *via* PCR with biotin and NTA modified primers is a convenient method to characterize most proteins for multiplexed SMFS without further modifications.

Analysis of the rupture force distribution reveals distinct differences between CBM3a WT and its Y67A mutant. These differences arise from the absence of the stabilizing tyrosine at position 67, potentially changing available binding sites of the mutant to the cellulose surface. The fact that no rupture forces greater than 25 pN were measured for Y67A at 1 pN/s could be related to the difference in sample size (N=259 vs. N=138 for WT and Y67A, respectively) as the tail of larger rupture forces at 0.1 pN/s is similar for WT and Y67A (N=161 vs N=159 for WT and Y67A, respectively). A similar shape of rupture force distributions was observed in previous AFM-based studies for CBM3a (27) and CBM1 (24, 56) but previous AFM analysis showed a more Gaussian-like distribution for CBM3a (28), (25). King et al (27) showed that specific binding of CBM3a can be blocked with the addition of NCC and restored by washing the CBM-functionalized AFM tip with an excess of water. There, both the initial and restored rupture force distributions displayed tailing, suggesting that non-specific binding was not the reason for the observation of higher rupture forces.

Surface diffusion of cellulases on crystalline cellulose was experimentally verified, although the extent of surface diffusion was minor compared to dynamic binding and unbinding to the substrate (57), (58). To date, no active motility or processive motion has been observed experimentally for CBMs without being tethered to a CD. However, a computational study of CBM1 from *T. reesei* revealed that CBM1 can diffuse from the hydrophilic to the hydrophobic surface of a cellulose I crystal during which multiple local energy minima with distinct orientations were sampled (50). Similarly, it has been shown that CBM1 can bind in a non-canonical orientation when binding to cellulose III (29), which further indicates that type-A CBMs potentially display a much larger range of binding orientations on crystalline cellulose surfaces. Single-molecule imaging revealed that CBM1 exhibits distinct surface binding events (22), which could be correlated to distinct regions of crystalline cellulose, and such binding modes may be found in structurally similar type-A CBMs such as CBM3a. When fused to an endoglucanase, CBM3a occupies more binding sites on crystalline cellulose compared to CBM1 fused to the same CD, further suggesting the presence of specific binding sites accessible to different type-A CBMs (59). The Y67A mutation is located at the edge of the binding site of CBM3a, thus effectively reducing the total planar binding motif. Consequently, binding orientations or binding sites which were determined or stabilized by the Y67-substrate interaction may no longer be favorable in the absence of this residue. The observation of only one rupture force peak and a slight change in the magnitude of binding forces for Y67A compared to the WT indicates the recognition of different binding sites on crystalline cellulose. This hypothesis is further supported by the reduction in total available binding sites by 1.8-fold despite similar unbinding rates as determined by QCM-D.

The tailing of the rupture force distributions towards larger rupture forces may also be correlated with the naturally evolved functional role of the CBM in the cellulosome. As cellulosomal microbes colonize cellulosic substrates, they are subjected to high interfacial shear forces, for example in the gut-intestine of higher organisms or sewage sludge (60). The main cellulosomal scaffold protein cohesin, is relatively stable and unfolds only under forces > 140 pN (61) (62), leaving the interdomain CBM mostly intact (63). This suggests that CBMs may have evolved to remain bound to cellulose during elevated levels of mechanical stress, but also remain flexible enough for the cellulosome to adopt to different conformations during hydrolysis (5) (64), which is reflected in our observed broad rupture force distribution. Nevertheless, multivalency in the form of multiple CDs interacting with the substrate may be as important in withstanding mechanical stress by the cellulosome but this has not yet been well characterized.

The multimodal rupture force distribution of the CBM3a WT suggests the existence of specific binding sites on the crystalline cellulose surface. To further validate this hypothesis and understand the role of each binding residue in the recognition of and dissociation from the substrate, rupture force measurements of mutants (such as H57 or W118 mutated to alanine), are suggested in future studies. While the application of the DHS model (49) for CBM3a WT yielded an unbinding rate comparable with previous single-molecule results, it still failed to accurately predict the unbinding rate of the Y67A mutant as well as describe the broad rupture force distribution with the obtained fit parameters. This issue might be resolved if bond-lifetime measurements of CBMs are carried out in force-clamp mode rather than transforming rupture force histograms to bond-lifetime data. Understanding the influence of each binding residue on the binding and unbinding rate will pave the way for specific rational engineering approaches to fine-tune CBM-substrate interaction for optimized catalytic activity and other designed functions.

## Materials and Methods

### Chemicals and substrates

Unless otherwise mentioned, all reagents were either purchased from VWR International, Fisher Scientific, USA, or Sigma-Aldrich, USA. Streptavidin-coated polystyrene particles (SVP30) with a nominal diameter of 3.11 μm were purchased from Spherotech Inc, USA. Amino-functionalized beads (01-01-503) with a nominal diameter of 5 μm were purchased from Micromod Partikeltechnologie GmbH and used as fiducial beads to account for drift during AFS assays. Sulfuric acid-hydrolyzed nanocrystalline cellulose was kindly donated by Richard Reiner at the USDA Forest Product Laboratory (65).

### DNA tethers

Linear double-stranded DNA tethers were synthesized in one step by PCR using the pEC-GFP-CBM3a plasmid as a template and 5’ modified primers. The biotin-modified primer (forward primer, 5’-biotin-C6-GGCGATCGCCTGGAAGTA) was purchased from Integrated DNA Technologies, Inc. USA. The NTA modified primer (backward primer, 5’-NTA-SS-C6-TCCAAAGGTGAAGAACTGTTCACC) was purchased from Gene Link, Inc. USA. The whole plasmid (5.4 kb) was amplified, then purified using the PCR Clean-up kit (IBI Scientific USA) resulting in a linear DNA tether of ~1.8 μm length with one modification on each end of the DNA. Amplification and product purity was verified by gel electrophoresis. In addition, a linear DNA tether of the same length was amplified using a digoxigenin-modified primer instead of NTA (5’-DIG-NHS-TCCAAAGGTGAAGAACTGTTCACC, Integrated DNA Technologies, Inc. USA) to bind to anti-digoxigenin Fab fragment antibodies (11214667001, Roche).

### Proteins

His_8_-GFP-CBM3a wild type (WT) and its Y67A mutant were expressed and purified as described previously (14).

### Buffers

All AFS experiments were carried out in working buffer (WB) containing 10 mM phosphate buffer at pH 7.4 supplemented with 0.31 mg/ml BSA and casein and 0.19 mg/ml Pluronic F-127, respectively. In addition, two blocking buffers were used to passivate the surface before the experiment. Buffer B1 consists of 10 mM phosphate buffer supplemented with 2.5 mg/ml BSA and casein. Buffer B2 consists of 10 mM phosphate buffer supplemented with 2.2 mg/ml BSA and casein and 5.6 mg/ml Pluronic F-127 respectively. All buffers were degassed in a vacuum (−90 kPa) for 30 minutes.

### QCM-D experiments

Quartz Crystal Microbalance with dissipation experiments were carried out and analyzed as described previously (14) except for using 10 mM phosphate buffer at pH 7.4 and sulfuric acid-derived NCC.

### Cellulose film preparation and AFS chip cleaning

The microfluidic chips used in the AFS are custom designed by LUMICKS B.V., the Netherlands for re-use. Therefore, a reliable protocol for the immobilization and removal of NCC needed to be established. To obtain a stable cellulose film, a multilayer deposition process (44) using an automated microfluidic control system was employed from Elvesys S.A.S, France. The system consists of a microfluidic controller (OB1, driven by compressed nitrogen), a 10-port distribution valve (MUX-D), pressurized fluid reservoirs (2-50 ml), and a manifold. To avoid potential damage to the 10-port valve when in contact with NCC, the valve was used to direct the pressurized nitrogen to the correct reservoir instead of directly controlling the liquid streams. Due to this configuration, installing check-valves on each line was necessary prior to entering the manifold to avoid backflow and cross-contamination between reservoirs. The flowsheet of the setup is shown in Figure 2 (panel A and B) and the detailed part list can be found in SI Appendix Table S4. The microfluidic resistance of the setup including the AFS chip was determined to be 3 *μl* / (*min* * *mbar*) and the volume flown through the chip was calculated based on the set pressure and duration. First, the cleaned chip was rinsed with 2 ml DI water, followed by flushing through 200 μl 0.05% (w/v) PLL and incubation for 1 minute. Next, the chip was rinsed with 1ml DI water and blow-dried for 1 minute. 200 μl of NCC at a concentration of 0.5% (w/v) was incubated for 1 minute, followed by 1ml water rinse and drying for 1 minute. The deposition of PLL and NCC was repeated four more times. Following the final NCC layer deposition, the chip was blow-dried for 20 minutes. Finally, the chip was disassembled and the bottom part which includes the flow cell was placed in an oven at 50°C to dry up overnight.

To confirm cellulose deposition using AFM, flow cells of the same channel geometry as the AFS chips were prepared by cutting the channel from *Parafilm*^®^ and fixing it between two microscope slides. Holes were drilled in one slide to connect 1/16” OD (1/32” ID) PTFE tubing. After assembly, the multilayer deposition process described above was employed manually. The slides were taken apart and dried up overnight at 50°C and stored in a desiccator until AFM imaging. The deposited NCC samples were visualized from the randomly selected area by an AFM (NX-10, Park systems). The AFM was used in non-contact mode operation with a scan size between 2×2 μm and 5×5 μm, 0.3 Hz scan rate, and 11.1 nm set point with the non-contact mode AFM tip (SSS-NCHR, Park systems). The AFM images were analyzed using XEI software (Park systems).

To directly verify the deposition of NCC inside the AFS chip, the fluorescence intensity of GFP-CBM3a bound to NCC was measured. All experiments were carried in at least triplicates. The chip was first rinsed with 500 μl DI water and 500 μl phosphate buffer followed by 15-minute passivation of the surface in B1 and B2 buffer, respectively. GFP-CBM3a WT was diluted in WB to a concentration of 1 μM and incubated for 5 minutes, followed by rinsing 1ml of WB. The fluorescence images were taken with a CMOS camera (Kiralux, Thorlabs Inc. USA) using μManager (66) on an inverted fluorescence microscope (Olympus IX 71) equipped with the necessary filters to enable GFP fluorescence. Control experiments on bare glass and PLL treated chip surfaces were performed to estimate the degree of non-specific binding of GFP-CBM3a. All images were corrected for background and shading (67).

The NCC was removed by incubating piranha solution (7:3 concentrated H_2_SO_4_:30% H_2_O_2_, v/v) two times for 15-30 minutes at 50°C with 500 μl DI water rinses in between. The next step in the cleaning procedure involved incubation of 1 M NaOH for 1-12 hours at room temperature followed by incubation of piranha solution for 15-30 minutes at 50°C, rinsing with 5 ml DI water, and drying. If the AFS chips were used for single-molecule experiments, 5 μm NH_2_-functionalized beads (to serve as fiducial beads) were diluted ~1:1000 in 0.01 M HCl and dried up inside the chip overnight at 50°C before the chip was functionalized with NCC.

### Tethered bead preparation for single-molecule force spectroscopy

Single-molecule experiments were carried out on a G1 AFS instrument with G2 AFS chips provided by LUMICKS B.V. After immobilizing NCC, the AFS chip was rinsed with 500 μl DI water and 500 μl phosphate buffer phosphate. Next, the surface was passivated with B1 and B2 buffer for 15 minutes each and rinsed with WB. The NTA-DNA tethers were diluted to 6 pM in WB containing 6 nM NiCl2. The bead-DNA-CBM construct was prepared in a two-step procedure. First, 15μl streptavidin-coated beads and Ni-NTA-DNA tethers were mixed to yield less than 1 DNA tether per bead and incubated on a rotisserie for 30 minutes. Details about the specificity of the Ni-NTA moiety for His-tagged CBMs can be found in **SI Appending Fig. S5** and the determination of the overall binding efficiency of DNA tethers to the beads is described in **SI Appendix**. The functionalized beads were washed twice by spinning down, removing the supernatant, and resuspending in 100 μl WB. GFP-CBM3a WT or Y67A mutant were diluted to 2 nM in WB. The washed and DNA functionalized bead pellet is resuspended in 20 μl of either WT or Y67A solution (resulting in a >1000x molar excess of CBM with respect to DNA) and placed on the rotisserie for 30 minutes. Next, the beads were washed twice in WB to remove any unbound CBM and resuspended in 20 μl WB or B2 if a high non-specific bead binding was observed during SFMS experiments. There was no significant difference in the partition coefficient between WB and B2 for WT (p=0.68, df=7) and Y67A (p=0.49, df=7) mutant. Refer to SI Appendix for information about the experimental setup **SI Appendix Fig. S6** for binding data. The CBM-DNA-bead construct was flushed through the AFS chip and incubated for 30 minutes. Non-bound beads were subsequently washed out with WB at a flow rate of 2 μl/min using a syringe pump (New Era Pump Systems Inc., USA). A small force of ~0.2-0.5 pN was applied to speed up the flushing step. For illustration, a schematic of the single-molecule setup is shown in **Figure 1** After measuring the rupture forces, the chip was rinsed with 100 μl WB, and the next CBM-DNA-bead sample was inserted.

To verify that the amplified DNA tethers are 1.8 μm in length, anti-digoxigenin fab fragments dissolved in PBS (20μg/ml) were non-specifically bound to the AFS glass surface for 20 minutes, followed by the same passivation procedure as outlined above. The DNA tethers in this experiment were functionalized with digoxigenin instead of NTA and prepared in one step since no protein needed to be attached to the bead, and a schematic is shown in **Figure 1** for illustration. The DNA-to-bead ratio was between 5-10 to ensure a sufficient yield of single-molecule tethers. DNA-functionalized beads were incubated on the surface between 10-30 minutes, the flushing process, bead tracking, and analysis procedure were identical to CBM-tethered bead experiments.

### Bead tracking, force ramp application, and determination of rupture forces

Tracking and analysis of the beads were accomplished using the software package provided by LUMICKS, with slight modifications to allow efficient export of rupture forces and associated tethers statistics as well as force-distance curves to a spreadsheet. The procedure for identifying a single-molecule tether, force calibration, and rupture force determination is described in detail elsewhere (30). The beads were tracked at 20 Hz using a 4x magnification objective. The trajectory of the beads without applied force was monitored for 8-10 minutes to determine the point of surface attachment (anchor point). Next, the force on each bead was calibrated by applying a constant amplitude for 2-4 minutes. Typically, 2-3 different amplitude values were used to build the calibration curve between the applied amplitude and effective force on each bead. Single-molecule tethers were identified by the root-mean-square fluctuation (*RMS*) and symmetrical motion (*Sym*) of the bead around the anchor point during the time frame for anchor point determination. For the CBM-cellulose experiment, values of single-molecule tethers for *RMS* and *Sym* are in the range between 850-1200 nm and 1.0-1.3 respectively. During force calibration, the diffusion coefficient of the bead and the force were used as fit parameters. This diffusion coefficient was compared to the diffusion coefficient determined by the Stokes-Einstein relation and was in the range between 0.8-1.2 for single tethers. The force obtained during force calibration was used to estimate the theoretical extension of DNA using the extensible WLC model (46). This extension was compared to the measured length during that force calibration point to yield the dimensionless length *l_fc_* and was expected to be close to 1 for single tethers. Next, a linear force ramp of either 0.1 or 1 pN/s was applied. Rupture forces were determined through the software by finding the time frame at which the z-position of the bead was outside the interval covered by the lookup-table (LUT) value (30). An example time trace of a typical rupture force measurement is shown in **SI Appendix Fig. S7**. Each trace and force-extension curve (FD) during force ramp application was inspected manually to determine the rupture force accurately.

### Analysis of rupture forces

Further evaluation of traces as well as data analysis was carried out by a custom-written MATLAB^®^ script (available on request). For each known single-molecule trace, several indicators such as *RMS*, *Sym*, *l_fc_*, rupture force and loading rate, along with the obtained force-distance (FD) curve during force ramp application for each trace, were imported into MATLAB^®^. To each FD curve, the dimensionless contour length *l_c_* of the WLC model based on the expected contour length of 1800 nm was fitted using a persistence length of *l_p_*=42 nm and stretch modulus *S*=1300 pN (68). This fitted length (determined during the force ramp) was compared to the dimensionless length during force calibration *l_fc_*, and only traces close to 1 were further analyzed. Traces in which the rupture force or loading rate was 3 standard deviations away from the sample mean were examined manually and discarded if the FD curve or any other mentioned statistics indicate that the trace did not originate from a single-tethered bead. To ensure that no bias was introduced by removing traces, the remaining data was subjected to a Pearson and Spearman correlation coefficient test between the obtained rupture force and *RMS*, *Sym* and *l_fc_* respectively. Next, the obtained rupture force histograms were converted to force dependent bond-lifetime data and analyzed using the procedure outlined by Dudko *et al*. (49) to obtain the bond-lifetime in the absence of force which is briefly described below. The rupture force histograms were converted to force-dependent bond-lifetime data using Equation 1:

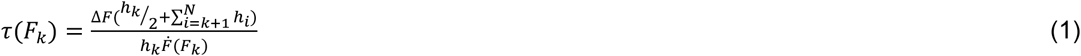

Where *τ*(*F_k_*) and 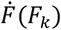 are the average bond-lifetime and loading rate at the k^th^ bin and 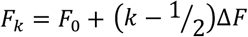. The rupture force histogram is composed of *N* bins of width Δ*F* starting from *F_0_* and ending at *F_0_* + *N*Δ*F*. The number of counts in the i^th^ bin is *C_i_* and the height of each bin can be calculated as 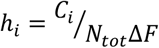 where *N_tot_* is the total number of counts.

The force-dependent bond-lifetime *τ*(*F*) is described using Equation 2:

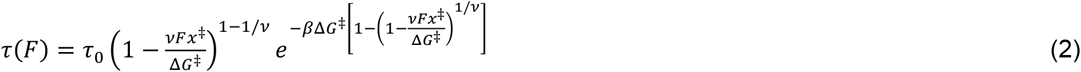

Where 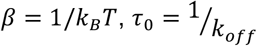 is the bond-lifetime (or inverse of the unbinding rate *k_off_*), x^‡^ is the transition state distance and Δ*G*^‡^ the apparent free energy of activation in the absence of the external force. The shape factor *v*=1/2 or 2/3 describes the underlying free-energy profile as cusp or linear-cubic, respectively.

The distribution of rupture forces is described by Equation 3:

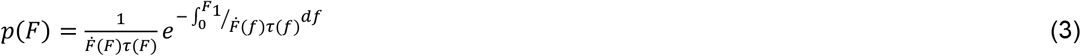

## Acknowledgments

Prof. Shishir P.S. Chundawat acknowledges support from the NSF (CBET Award 1846797), Rutgers Aresty Research Center, and Rutgers SOE Startup Funds. John M. Yarbrough was supported by the DOE, Office of Energy Efficiency and Renewable Energy under Agreement No. 28598. Prof. Ki-Bum Lee acknowledges partial support from the NSF (CBET-1803517), the New Jersey Commission on Spinal Cord (CSCR17IRG010; CSCR16ERG019), and NIH R01 (1R01DC016612, 3R01DC016612-01S1, and 5R01DC016612-02S1). Markus Hackl would like to thank Bhargava Nemmaru for help with the QCM-D data analysis. Figure 1-A,C,D, Figure S5-A, and Figure S7-A were created with BioRender.com.

## References

1. S. P. S. Chundawat, G. T. Beckham, M. E. Himmel, B. E. Dale, Deconstruction of Lignocellulosic Biomass to Fuels and Chemicals. Annu. Rev. Chem. Biomol. Eng. 2, 121 – 145 (2011).

2. M. E. Himmel, et al., Biomass recalcitrance: Engineering plants and enzymes for biofuels production. Science (80−.). 315, 804–807 (2007).

3. C. M. Payne, et al., Fungal cellulases. Chem. Rev. 115, 1308–1448 (2015).

4. C. M. G. A. Fontes, H. J. Gilbert, Cellulosomes: Highly efficient nanomachines designed to deconstruct plant cell wall complex carbohydrates. Annu. Rev. Biochem. 79, 655–681 (2010).

5. M. Eibinger, T. Ganner, H. Plank, B. Nidetzky, A Biological Nanomachine at Work: Watching the Cellulosome Degrade Crystalline Cellulose. ACS Cent. Sci. 6, 739–746 (2020).

6. E. A. Bayer, H. Chanzy, R. Lamed, Y. Shoham, Cellulose, cellulases and cellulosomes. Curr. Opin. Struct. Biol. 8, 548–557 (1998).

7. R. H. Doi, A. Kosugi, Cellulosomes: Plant-cell-wall-degrading enzyme complexes. Nat. Rev. Microbiol. 2, 541–551 (2004).

8. K. Igarashi, et al., Traffic Jams Reduce Hydrolytic Efficiency of Cellulase on Cellulose Surface. Science (80−.). 333, 1279–1282 (2011).

9. Y. Zhang, M. Zhang, R. Alexander Reese, H. Zhang, B. Xu, Real-time single molecular study of a pretreated cellulose hydrolysis mode and individual enzyme movement. Biotechnol. Biofuels 9, 85 (2016).

10. Y. Shibafuji, et al., Single-molecule imaging analysis of elementary reaction steps of trichoderma reesei cellobiohydrolase i (Cel7A) hydrolyzing crystalline cellulose lα and IIII. J. Biol. Chem. 289, 14056–14065 (2014).

11. B. W. McLean, et al., Analysis of binding of the family 2a carbohydrate-binding module from Celludomonas fimi xylanase 10a to cellulose: Specificity and identification of functionally important amino acid residues. Protein Eng. 13, 801–809 (2000).

12. P. J. Kraulis, et al., Determination of the Three-Dimensional Solution Structure of the C-Terminal Domain of Cellobiohydrolase I from Trichoderma reesei. A Study Using Nuclear Magnetic Resonance and Hybrid Distance Geometry-Dynamical Simulated Annealing. Biochemistry 28, 7241–7257 (1989).

13. J. Tormo, et al., Crystal structure of a bacterial family-III cellulose-binding domain: a general mechanism for attachment to cellulose. EMBO J. 15, 5739–5751 (1996).

14. B. Nemmaru, et al., Reduced type-A carbohydrate-binding module interactions to cellulose I leads to improved endocellulase activity. Biotechnol. Bioeng. 118, 1141–1151 (2021).

15. D. Gao, et al., Increased enzyme binding to substrate is not necessary for more efficient cellulose hydrolysis. Proc. Natl. Acad. Sci. U. S. A. 110, 10922–10927 (2013).

16. J. Kari, et al., Physical constraints and functional plasticity of cellulases. Nat. Commun. 12, 1–10 (2021).

17. D. W. Abbott, A. B. Boraston, Quantitative approaches to the analysis of carbohydrate-binding module function. Methods Enzymol. 510, 211–231 (2012).

18. A. L. Creagh, E. Ong, E. Jervis, D. G. Kilburn, C. A. Haynes, Binding of the cellulose-binding domain of exoglucanase Cex from Cellulomonas fimi to insoluble microcrystalline cellulose is entropically driven. Proc. Natl. Acad. Sci. 93, 12229–12234 (1996).

19. Y. Zhang, et al., Interactions between type A carbohydrate binding modules and cellulose studied with a quartz crystal microbalance with dissipation monitoring. Cellulose 4 (2020).

20. J. Guo, J. M. Catchmark, Binding specificity and thermodynamics of cellulose-binding modules from trichoderma reesei Cel7A and Cel6A. Biomacromolecules 14, 1268–1277 (2013).

21. S. J. Charnock, et al., Promiscuity in ligand-binding: The three-dimensional structure of a Piromyces carbohydrate-binding module, CBM29-2, in complex with cello-and mannohexaose. Proc. Natl. Acad. Sci. U. S. A. 99, 14077–14082 (2002).

22. A. Nakamura, et al., Single-molecule imaging analysis of binding, processive movement, and dissociation of cellobiohydrolase trichoderma reesei Cel6A and its domains on crystalline cellulose. J. Biol. Chem. 291, 22404–22413 (2016).

23. S. K. Brady, S. Sreelatha, Y. Feng, S. P. S. Chundawat, M. J. Lang, Cellobiohydrolase 1 from Trichoderma reesei degrades cellulose in single cellobiose steps. Nat. Commun. 6, 10149 (2015).

24. A. Griffo, et al., Binding Forces of Cellulose Binding Modules on Cellulosic Nanomaterials. Biomacromolecules 20, 769–777 (2019).

25. M. Zhang, S.-C. Wu, W. Zhou, B. Xu, Imaging and Measuring Single-Molecule Interaction between a Carbohydrate-Binding Module and Natural Plant Cell Wall Cellulose. J. Phys. Chem. B 116, 9949–9956 (2012).

26. M. Zhang, B. Wang, B. Xu, Mapping single molecular binding kinetics of carbohydrate-binding module with crystalline cellulose by atomic force microscopy recognition imaging. J. Phys. Chem. B 118, 6714–6720 (2014).

27. J. R. King, C. M. Bowers, E. J. Toone, Specific binding at the cellulose binding module-cellulose interface observed by force spectroscopy. Langmuir 31, 3431–3440 (2015).

28. M. Zhang, B. Wang, B. Xu, Measurements of single molecular affinity interactions between carbohydrate-binding modules and crystalline cellulose fibrils. Phys. Chem. Chem. Phys. 15, 6508–6515 (2013).

29. S. P. S. Chundawat, et al., Molecular origins of reduced activity and binding commitment of processive cellulases and associated carbohydrate-binding proteins to cellulose III. J. Biol. Chem. 296, 100431 (2021).

30. G. Sitters, et al., Acoustic force spectroscopy. Nat. Methods 12, 47–50 (2014).

31. D. Kamsma, R. Creyghton, G. Sitters, G. J. L. Wuite, E. J. G. Peterman, Tuning the Music: Acoustic Force Spectroscopy (AFS) 2.0. Methods 105, 26–33 (2016).

32. F. Ritort, Single-molecule experiments in biological physics: Methods and applications. J. Phys. Condens. Matter 18 (2006).

33. E. Evans, Energy landscapes of biomolecular adhesion and receptor anchoring at interfaces explored with dynamic force spectroscopy. Faraday Discuss., 1–16 (1998).

34. F. Rico, V. T. Moy, Energy landscape roughness of the streptavidin–biotin interaction. J. Mol. Recognit. 20, 495–501 (2007).

35. R. P. Goodman, et al., A facile method for reversibly linking a recombinant protein to DNA. ChemBioChem 10, 1551–1557 (2009).

36. G. D. Meredith, H. Y. Wu, N. L. Allbritton, Targeted protein functionalization using his-tags. Bioconjug. Chem. 15, 969–982 (2004).

37. J. Shimada, T. Maruyama, M. Kitaoka, N. Kamiya, M. Goto, DNA-enzyme conjugate with a weak inhibitor that can specifically detect thrombin in a homogeneous medium. Anal. Biochem. 414, 103–108 (2011).

38. J. Shimada, et al., Conjugation of DNA with protein using His-tag chemistry and its application to the aptamer-based detection system. Biotechnol. Lett. 30, 2001–2006 (2008).

39. L. Schmitt, M. Ludwig, H. E. Gaub, R. Tampé, A metal-chelating microscopy tip as a new toolbox for single-molecule experiments by atomic force microscopy. Biophys. J. 78, 3275–3285 (2000).

40. C. Verbelen, H. J. Gruber, Y. F. Dufrêne, The NTA-His6 bond is strong enough for AFM single-molecular recognition studies. J. Mol. Recognit. 20, 490–494 (2007).

41. F. Kienberger, et al., Recognition Force Spectroscopy Studies of the NTA-His6 Bond. Single Mol. 1, 59–65 (2000).

42. M. Conti, G. Falini, B. Samorì, How strong is the coordination bond between a histidine tag and Ni-nitrilotriacetate? An experiment of mechanochemistry on single molecules. Angew. Chemie - Int. Ed. 39, 215–218 (2000).

43. R. W. Friddle, A. Noy, J. J. De Yoreo, Interpreting the widespread nonlinear force spectra of intermolecular bonds. Proc. Natl. Acad. Sci. 109, 13573–13578 (2012).

44. E. D. Cranston, D. G. Gray, Morphological and optical characterization of polyelectrolyte multilayers incorporating nanocrystalline cellulose. Biomacromolecules 7, 2522–2530 (2006).

45. E. Kontturi, et al., Cellulose nanocrystal submonolayers by spin coating. Langmuir 23, 9674–9680 (2007).

46. T. Odijk, Stiff Chains and Filaments under Tension. Macromolecules 28, 7016–7018 (1995).

47. D. Freedman, P. Diaconis, On the histogram as a density estimator:L2 theory. Zeitschrift für Wahrscheinlichkeitstheorie und Verwandte Gebiete 57, 453–476 (1981).

48. O. K. Dudko, G. Hummer, A. Szabo, Intrinsic rates and activation free energies from single-molecule pulling experiments. Phys. Rev. Lett. 96, 1–4 (2006).

49. O. K. Dudko, G. Hummer, A. Szabo, Theory, analysis, and interpretation of singlemolecule force spectroscopy experiments. Proc. Natl. Acad. Sci. U. S. A. 105, 15755–15760 (2008).

50. M. R. Nimlos, et al., Binding preferences, surface attachment, diffusivity, and orientation of a family 1 carbohydrate-binding module on cellulose. J. Biol. Chem. 287, 20603–20612 (2012).

51. C. Aulin, et al., Nanoscale Cellulose Films with Different Crystallinities and Mesostructures—Their Surface Properties and Interaction with Water. Langmuir 25, 7675–7685 (2009).

52. S. Ahola, J. Salmi, L.-S. Johansson, J. Laine, M. Österberg, Model Films from Native Cellulose Nanofibrils. Preparation, Swelling, and Surface Interactions. Biomacromolecules 9, 1273–1282 (2008).

53. R. J. Moon, A. Martini, J. Nairn, J. Simonsen, J. Youngblood, Cellulose nanomaterials review: structure, properties and nanocomposites. Chem. Soc. Rev. 40, 3941 (2011).

54. T. Abitbol, A. Palermo, J. M. Moran-Mirabal, E. D. Cranston, Fluorescent labeling and characterization of cellulose nanocrystals with varying charge contents. Biomacromolecules 14, 3278–3284 (2013).

55. Z. Huang, V. S. Raghuwanshi, G. Garnier, Functionality of immunoglobulin G and immunoglobulin M antibody physisorbed on cellulosic films. Front. Bioeng. Biotechnol. 5, 1–10 (2017).

56. B. Arslan, et al., The Effects of Noncellulosic Compounds on the Nanoscale Interaction Forces Measured between Carbohydrate-Binding Module and Lignocellulosic Biomass. Biomacromolecules 17, 1705–1715 (2016).

57. J. Moran-Mirabal, J. Bolewski, L. P. Walker, Thermobifida fusca Cellulases Exhibit Limited Surface Diffusion on Bacterial Micro-Crystalline Cellulose. Biotech Bioeng 110, 47–56 (2013).

58. E. J. Jervis, C. A. Haynes, D. G. Kilburn, Surface diffusion of cellulases and their isolated binding domains on cellulose. J. Biol. Chem. 272, 24016–24023 (1997).

59. G. Carrard, A. Koivula, H. Söderlund, P. Béguin, Cellulose-binding domains promote hydrolysis of different sites on crystalline cellulose. Proc. Natl. Acad. Sci. U. S. A. 97, 10342–10347 (2000).

60. L. Artzi, E. A. Bayer, S. Moraïs, Cellulosomes: Bacterial nanomachines for dismantling plant polysaccharides. Nat. Rev. Microbiol. 15, 83–95 (2017).

61. T. Verdorfer, et al., Combining in Vitro and in Silico Single-Molecule Force Spectroscopy to Characterize and Tune Cellulosomal Scaffoldin Mechanics. J. Am. Chem. Soc. 139, 17841–17852 (2017).

62. A. Galera-Prat, S. Moraïs, Y. Vazana, E. A. Bayer, M. Carrión-Vázquez, The cohesin module is a major determinant of cellulosome mechanical stability. J. Biol. Chem. 293, 7139–7147 (2018).

63. S. W. Stahl, et al., Single-molecule dissection of the high-affinity cohesin-dockerin complex. Proc. Natl. Acad. Sci. 109, 20431–20436 (2012).

64. J. J. Adams, et al., Insights into higher-order organization of the cellulosome revealed by a dissect-and-build approach: Crystal structure of interacting Clostridium thermocellum multimodular components. J. Mol. Biol. 396, 833–839 (2010).

65. R. S. Reiner, A. W. Rudie, “Process scale-up of cellulose nanocrystal production to 25 kg per batch at the Forest Products Laboratory” in Production and Applications of Cellulose Nanomaterials, (TAPPI Press, 2013), pp. 21–24.

66. A. Edelstein, N. Amodaj, K. Hoover, R. Vale, N. Stuurman, Computer control of microscopes using manager. Curr. Protoc. Mol. Biol., 1–17 (2010).

67. M. A. Model, J. K. Burkhardt, A standard for calibration and shading correction of a fluorescence microscope. Commun. Clin. Cytom. 46, 309–316 (2001).

68. M. D. Wang, H. Yin, R. Landick, J. Gelles, S. M. Block, Stretching DNA with optical tweezers. Biophys. J. 72, 1335–1346 (1997).

