## Supplementary Information for "Acoustic Force Spectroscopy Reveals Subtle Differences in Cellulose Unbinding Behavior of Carbohydrate-Binding Modules"

**This PDF file includes:**

Supplementary text
Figures S1 to S7
Tables S1 to S4
SI References

### Supplementary text

**Biotinylated DNA binding efficiency to streptavidin-coated beads.** The binding efficiency was determined using a supernatant assay (1) in which the concentration of unbound DNA was measured using the Quant-it PicoGreen dsDNA Assay kit (Thermo Fisher Scientific, USA). For each experiment, 70 µl of 3.11 µm diameter streptavidin-coated beads were washed in Tris-EDTA (TE) buffer at pH 7 containing 1mg/ml BSA and mixed in a PCR tube with 1.8 µm long biotin/digoxigenin- DNA tethers at a ratio of 100 tethers per bead. Control experiments with the same concentration of DNA but without beads were used to estimate the amount of non-specific binding of DNA to the PCR tube. The bead-DNA mixture was incubated on a rotor for 30 minutes and subsequently spun down to separate beads and supernatant. The supernatant (50 µl) was transferred to a clear-bottom 96-well plate and mixed with the fluorescent dye following the manufacturer's protocol. The fluorescence was measured at 480 nm excitation, 520 nm emission with a cut-off of 495 nm. The concentration of DNA was determined using a standard curve of λ-DNA supplied with the kit. The binding efficiency was calculated as $BE = 1 - \frac{c_S}{c_A}$, where $c_S$ is the concentration of DNA in the supernatant of the bead containing sample and $c_A$ the concentration of DNA in the supernatant of the control sample. The degree of non-specific binding of DNA was calculated as $NS = 1 - \frac{c_A}{c_T}$ where $c_T$ is the concentration of DNA initially added. The binding efficiency of 1.8 µm long biotin/digoxigenin- DNA tethers was determined to be 8.8 ± 0.5% (mean±SD), whereas the degree of non-specific binding was significantly higher with 20.2 ± 0.87%.

**NTA-Tether specificity for His-tagged proteins.** To confirm the specificity of the NTA -DNA tether for His-tagged proteins, a similar approach as described in (2) was performed. Instead of attaching the NTA-DNA via the biotin handle to streptavidin-coated microplates, streptavidin-coated beads (20 µl) were used with a DNA-to-bead ratio of 1000, 10,000, and 20,000. The respective amount of DNA was added to the beads with 100-1000x molar excess of $NiCl_2$ and incubated in WB on a rotisserie for 30 minutes. Next, the beads were washed twice with WB followed by resuspension in 20µl WB containing His-tagged CelE-CBM3a (3) at a concentration of 285 nM followed by incubation for 15 minutes. This resulted in a molar excess of protein between 10-200 with respect to DNA. To test whether the protein is immobilized to the beads via the His-tag, half of the bead samples were washed three times with WB, whereas the other half was washed three times with WB containing 500 mM imidazole to elute the protein from the Ni-NTA tethers. All samples were resuspended in 60 µl working buffer containing 2mM pNP-cellobiose (4-Nitrophenyl-β-D cellobioside, Carbosynth Ltd, USA) and incubated at 50°C for 24 hours with overhead mixing. A schematic of the experimental setup is shown in **Fig. S5**-A. The concentration of released pNP was determined by measuring the absorbance and 405nm and comparison to a standard curve. **Fig. S5**-B shows the hydrolysis of pNP-cellobiose as a function of the molar DNA-to-bead ratio (amount of CelE-CBM3a immobilized per bead). The conversion increases with increasing CelE-CBM3a density on the beads. Control experiments in which CelE-CBM3a-functionalized beads have been washed with imidazole show no significant conversion, thus verifying that the prepared DNA tether specifically binds to streptavidin-coated beads and His-tagged proteins.

**Pull-down (solid-state depletion) assay for CBM3a-wt and Y67A mutant on NCC.** Pull-down assays of CBM3a WT and Y67A were performed in WB and B2 to investigate the effect of blocking agents on the binding affinity. The assays were carried out in clear 96-round bottom well plates in triplicates. To each well, 10 µl of 1 mg/ml NCC for WT and 10 µl of 10mg/ml of NCC for Y67A were added, followed by the addition of 90 µl of protein at a concentration between 0.25 µM and 4 µM. Control experiments for non-specific binding to the wells without substrate at the same protein concentrations were prepared on the same plate. The plates were covered with parafilm and incubated on a thermomixer at room temperature for 3 hours and shaking at 500 rpm. After binding, 7 µl of 5 M NaCl were added to precipitate the NCC, and the plates were centrifuged for 15min at 3200 g. The supernatant was transferred to an opaque 96-well flat bottom plate and the fluorescence was measured at 488 nm excitation with 509 nm emission and a 495 nm cut-off. The fluorescence intensity was converted to protein concentration based on a standard curve which was prepared parallel to the experiment. The concentration data were converted to bound protein per g of NCC and fitted to the partition coefficient described in Equation S1.

$$B = \frac{n_{max}}{K_d} * F \qquad \text{(S1)}$$

Where $B$ represents the amount of bound protein per gram of NCC and $F$ stands for the free protein concentration. The ratio of the maximum number of binding sites, $n_{max}$, and dissociation constant, $K_d$ is referred to as the apparent partition coefficient and describes the distribution of protein between substrate and solution in equilibrium. **Fig. S6** shows the binding data of CBM3a WT and Y67A mutant in WB and B2 along with the fit Equation S1. The partition coefficient for CBM3a WT is 6.5 ± 0.4 (mean±SE) L/g and 6.4 ± 0.7 L/g for WB and B2 respectively and 0.73 ± 0.05 L/g and 0.69 ± 0.11 L/g for the Y67A mutant in WB and B2 respectively.

**Supplementary Figures**

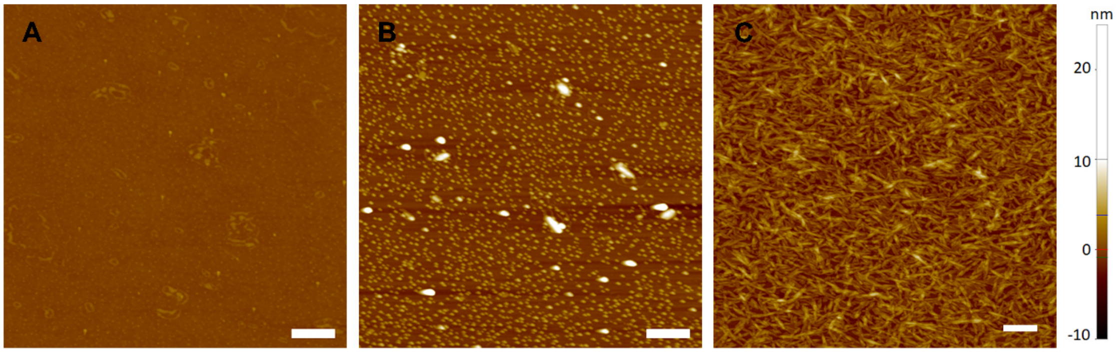


**Fig. S1. AFM images of glass slides at different stages of the multilayer deposition process.** A) bare surface after piranha treatment, B) poly-L-lysine (PLL) treated surface, C) 1x NCC layer on a PLL layer. The $R_a$ along randomly selected lines is 0.19 nm for the bare surface, 1.19 nm for the PLL treated surface, and 1.30 nm for one NCC layer on PLL. The scale bar is 500nm.

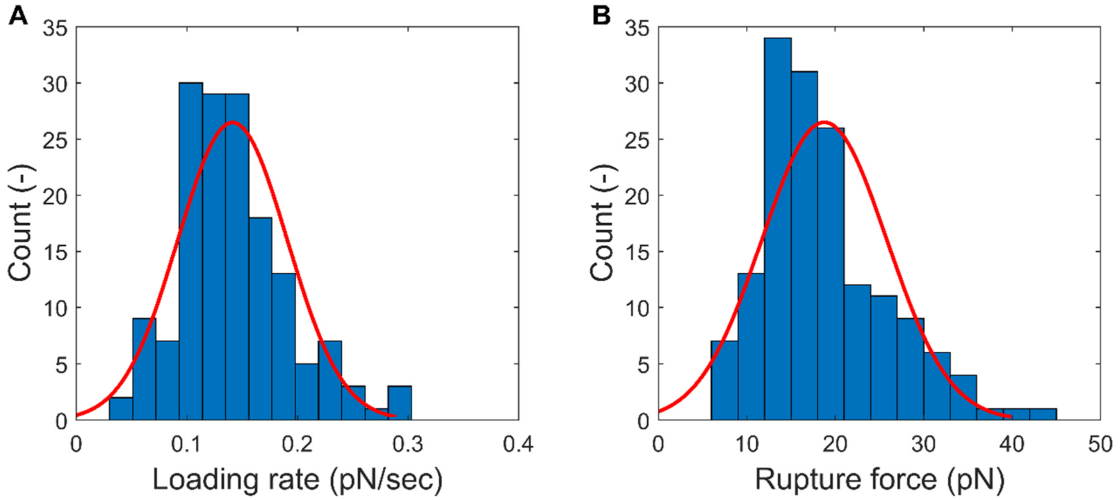


**Fig. S2. Digoxigenin-anti-digoxigenin (DIG-aDIG) unbinding forces measured using acoustic force spectroscopy.** Loading rate (A) and rupture force (B) histogram of DIG-aDIG (N=156). The fitted values (mean ± s.d.) are 0.14 ± 0.05 pN/s and 18.8 ± 7.0 pN respectively.

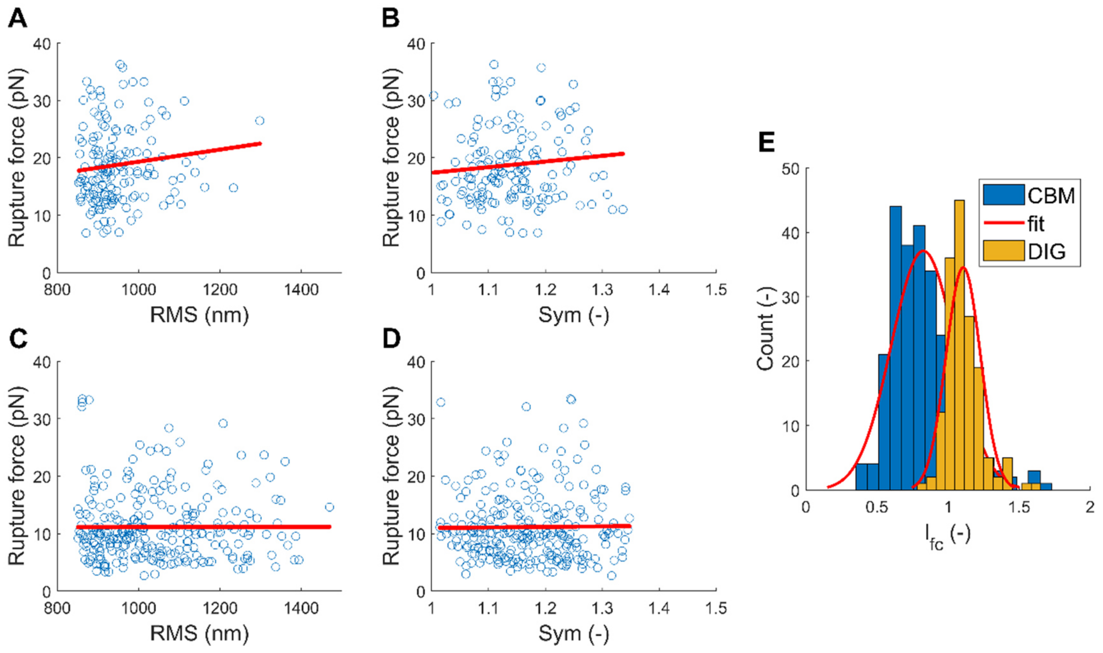


**Fig. S3. No correlation is found between the rupture force and $RMS$ or $Sym$, even though shorter tethers are observed on NCC surfaces.** A-B) Scatter plots of rupture force with $RMS$ and $Sym$ of DIG-aDIG rupture measurements and C-D) Scatter plots of rupture force with $RMS$ and $Sym$ of CBM3a WT- NCC rupture measurements at 1 pN/s. The red line indicates the best linear fit to the data. No significant correlation between rupture force and $RMS$ or $Sym$ is observed. Refer to **Table S1** and **Table S2** for correlation coefficients. E) Histogram of dimensionless length during force calibration ($l_{fc}$) for DIG-aDIG (yellow, N=156) and CBM3a WT-NCC (blue, N=259). The red line shows the normal distribution fit to each data set and mean±SD are 0.83 ± 0.23 and 1.1 ± 0.12 for CBM3a and DIG-aDIG, respectively. On average, a 25% reduction in measured length was observed for tethers on NCC surfaces**.**

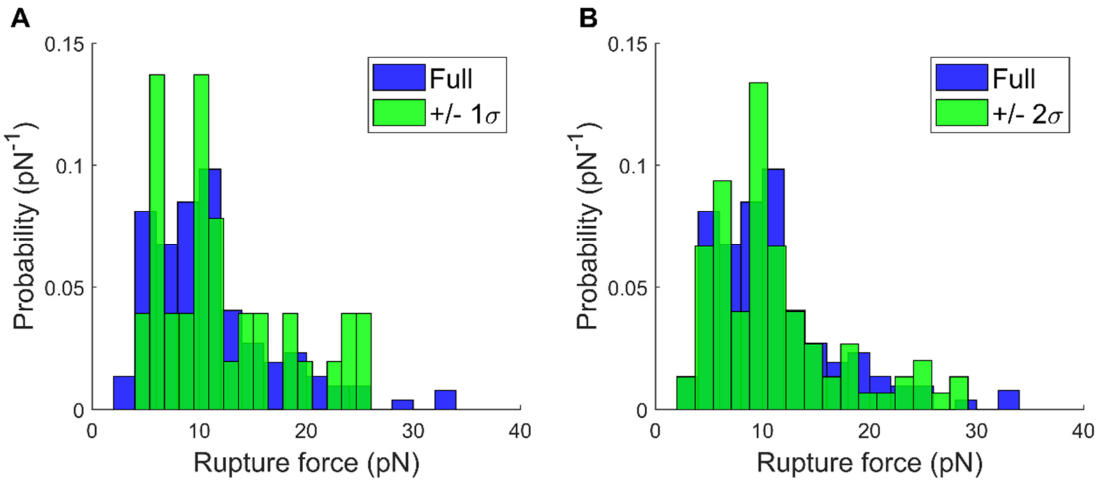


**Fig. S4. Comparison of entire rupture force histogram of CBM3a WT obtained at 1 pN/s (N=259) with reduced data sets.** The criteria for inclusion in the reduced data set is an expected length during force calibration ($l_{fc}$) identical to $l_{fc}$ obtained from DIG-aDIG experiments (μ ± σ=1.1 ± 0.12). The blue histogram represents the full data set, whereas the green histogram shows reduced data for A) μ ± σ (N=37) and B) μ ± 2σ (N=88). Qualitatively, the same shape of the histogram is obtained in both cases, thus indicating that shorter than expected DNA tethers are not affecting the rupture force measurement.

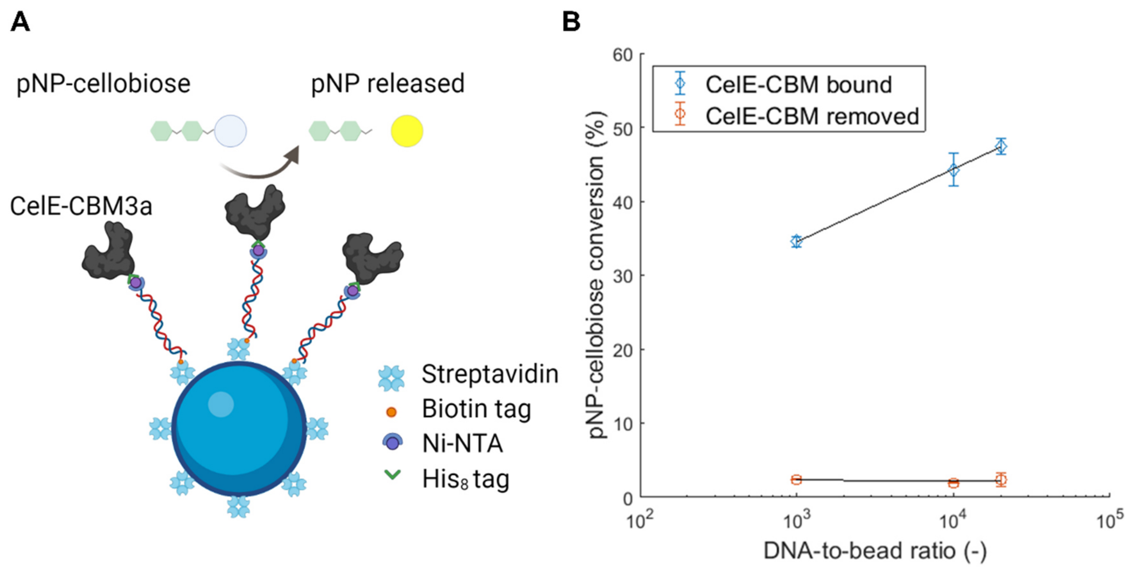


**Fig. S5. Validation of binding specificity of Ni-NTA modified DNA tethers to His-tagged proteins.** A) Schematic of the experimental setup. The DNA is connected to the bead by the streptavidin-biotin bond, whereas CelE-CBM3a is bound to the DNA by the Ni-NTA-Histidine bond. CelE-CBM3a hydrolyses the pNP-cellobiose to release pNP, which can be quantified by measuring the absorbance at 405 nm. B) Conversion of pNP-cellobiose as a function of molar DNA-to-bead ratio shows that samples washed with imidazole lose all the immobilized CelE and result in no significant conversion. Error bars represent the standard deviation of 3 experimental replicates. The line is the best logarithmic fit as a guide.

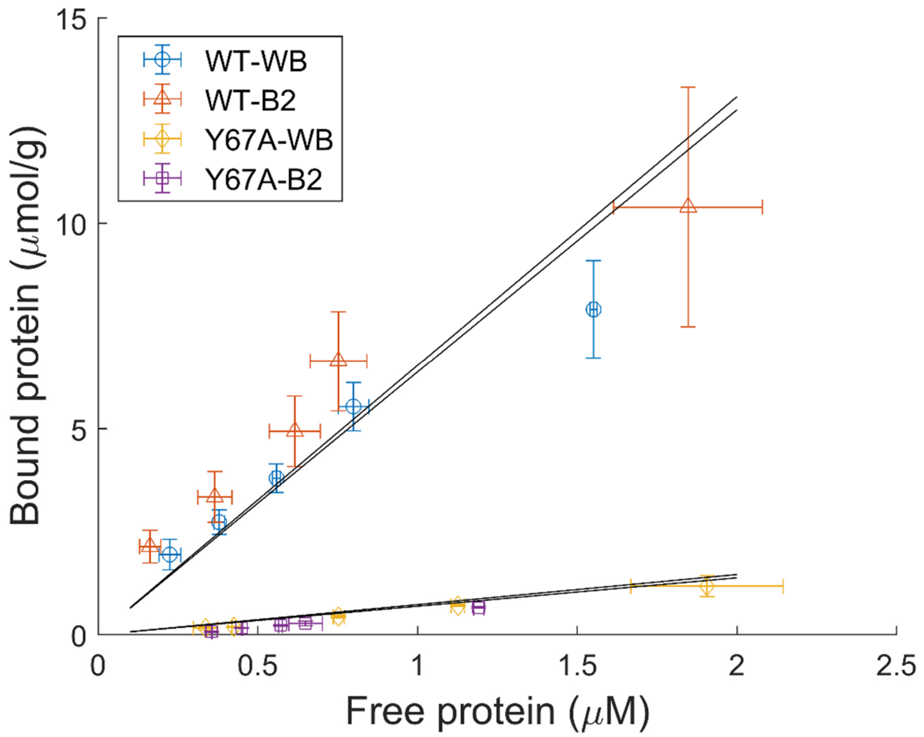


**Fig. S6. Bulk ensemble binding data of CBM3a WT and Y67a mutant to nanocrystalline cellulose.** Binding data of CBM3a WT in WB (o), B2 (Δ) and Y67A mutant in WB (◊) and B2 (□). Error bars indicate the standard deviation of three experiments. Solid lines represent the fit of equation S1 to the data to obtain the partition coefficient.

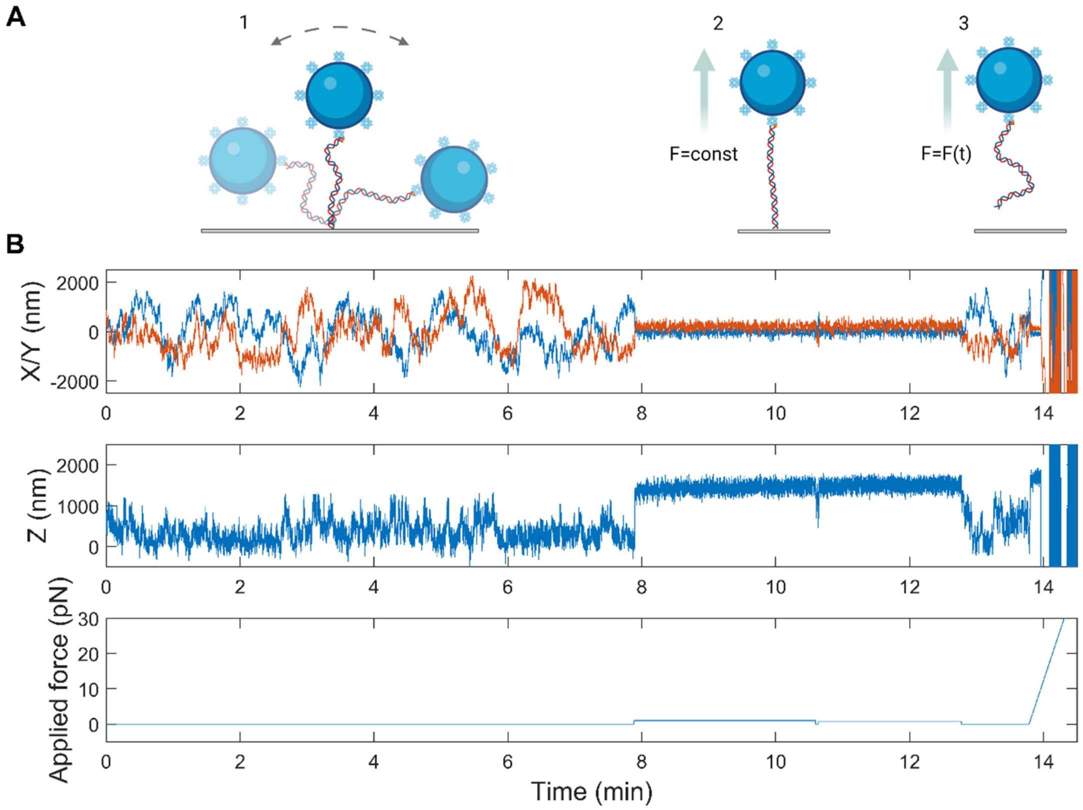


**Fig. S7. Overview of a typical AFS single-molecule rupture force assay.** A) Sketch of a tethered particle during anchor point determination (1), force calibration (2), and rupture during the force ramp (3). B) Example of a single-molecule trace recorded with the AFS. The anchor point was determined during the first 8 minutes at which the $RMS$ and $Sym$ values were calculated. Between minutes 8 and 12.5, the force was calibrated, and the measured extension was compared to the theoretical extension following the WLC model to yield $l_{fc}$. Finally, a linear force ramp was applied until the bond ruptured at approximately 14 minutes.

**Supplementary Tables**

**Table S1.** Pearson correlation coefficient with p-values in parenthesis tested between the measured rupture force and $RMS$, $Sym$ and $l_{fc}$ respectively. No significant correlation is observed except between rupture force and $Sym$ for Y67A at 0.1pN/s (p=0.043).

| | $\boldsymbol{RMS}$ | $\boldsymbol{Sym}$ | $\boldsymbol{l_{fc}}$ |
|---|---|---|---|
| **DIG-aDIG** | 0.1163 (0.148) | 0.0968 (0.229) | 0.0672 (0.404) |
| **WT 1pN/s** | 0.0024 (0.969) | 0.0126 (0.840) | 0.0109 (0.861) |
| **WT 0.1 pN/s** | 0.0480 (0.545) | 0.0382 (0.630) | -0.0128 (0.872) |
| **Y67A 1pN/s** | -0.083 (0.334) | -0.0741 (0.388) | -0.0268 (0.755) |
| **Y67A 0.1pN/s** | -0.0498 (0.533) | 0.1609 (0.043) | -0.0214 (0.789) |

**Table S2.** Spearman correlation coefficient with p-value in parenthesis tested between the measured rupture force and $RMS$, $Sym$ and $l_{fc}$ respectively. No significant correlation is observed except between rupture force and $Sym$ for Y67A at 0.1pN/s.

| | $\boldsymbol{RMS}$ | $\boldsymbol{Sym}$ | $\boldsymbol{l_{fc}}$ |
|---|---|---|---|
| **DIG-aDIG** | 0.1081 (0.179) | 0.1102 (0.171) | 0.0582 (0.471) |
| **WT 1pN/s** | 0.0217 (0.728) | -0.0158 (0.780) | -0.0004 (0.995) |
| **WT 0.1 pN/s** | 0.0329 (0.679) | 0.0274 (0.730) | -0.0185 (0.816) |
| **Y67A 1pN/s** | -0.1073 (0.210) | -0.0547 (0.524) | -0.0664 (0.439) |
| **Y67A 0.1pN/s** | -0.0229 (.0774) | 0.1497 (0.060) | -0.0074 (0.926) |

**Table S3.** Mean and standard deviation of double normal fit of the rupture force distribution for CBM3a-wt and Y67A mutant shown in Fig 4 B-C in the main manuscript. $p(\mu_1)$ denotes the fraction of the first peak.

| | | $p(\mu_1)$ (-) | $\mu_1$ (pN) | $\sigma_1$(pN) | $\mu_2$(pN) | $\sigma_2$(pN) |
|---|---|---|---|---|---|---|
| **WT** | 1 pN/s | 0.71 | 8.52 | 2.86 | 17.54 | 6.44 |
| | 0.1 pN/s | 0.37 | 3.50 | 0.69 | 7.06 | 2.66 |
| **Y67A** | 1 pN/s | 0.64 | 7.90 | 2.31 | 14.90 | 4.29 |
| | 0.1 pN/s | 0.67 | 4.52 | 1.13 | 9.08 | 3.67 |

**Table S4.** Parts list for the microfluidic setup shown in Figure 2-B in the main text.

| Item Description | QTY | Part Number | Vendor |
|---|---|---|---|
| Pressure Controller | 1 | OB1-MKIII+ | Elvesys S.A.S. |
| 10-way Valve | 1 | MUX-D | Elvesys S.A.S. |
| 50ml Fluid Reservoirs | 1 | KRM1 | Elvesys S.A.S. |
| 2ml Fluid Reservoirs | 2 | KRXS-V2 | Elvesys S.A.S. |
| PTFE Tubing 1/16” OD (1/32” ID) | 8m | KFSPPI | Elvesys S.A.S. |
| Low -Pressure Manifold Assembly, 6 port | 1 | P-152 | Idex Corporation |
| Adapter, Male Luer Lock x Female ¼-28 Flat Bottom | 4 | P-675 | Idex Corporation |
| Adapter, Female Luer x Male ¼-28 Flat Bottom | 4 | P-624 | Idex Corporation |
| Check Valve, Female Luer x Male Luer Lock | 4 | 30505-92 | Masterflex |

### SI References


1. E. W. A. Visser, L. J. Van Ijzendoorn, M. W. J. Prins, Particle Motion Analysis Reveals Nanoscale Bond Characteristics and Enhances Dynamic Range for Biosensing. *ACS Nano* **10**, 3093–3101 (2016).
2. J. Shimada, *et al.*, Conjugation of DNA with protein using His-tag chemistry and its application to the aptamer-based detection system. *Biotechnol. Lett.* **30**, 2001–2006 (2008).
3. C. K. Bandi, A. Goncalves, S. V. Pingali, S. P. S. Chundawat, Carbohydrate-binding domains facilitate efficient oligosaccharides synthesis by enhancing mutant catalytic domain transglycosylation activity. *Biotechnol. Bioeng.* **117**, 2944–2956 (2020).
